# River network rearrangements promote speciation in lowland Amazonian birds

**DOI:** 10.1101/2021.11.15.468717

**Authors:** Lukas J. Musher, Melina Giakoumis, James Albert, Glaucia Del-Rio, Marco Rego, Gregory Thom, Alexandre Aleixo, Camila C. Ribas, Robb T. Brumfield, Brian Tilston Smith, Joel Cracraft

## Abstract

Large Amazonian rivers impede dispersal for many species, but lowland river networks frequently rearrange, thereby altering the location and effectiveness of river-barriers through time. These rearrangements may promote biotic diversification by facilitating episodic allopatry and secondary contact among populations. We sequenced genome-wide markers to evaluate histories of divergence and introgression in six Amazonian avian species-complexes. We first tested the assumption that rivers are barriers for these taxa and found that even relatively small rivers facilitate divergence. We then tested whether species diverged with gene flow and recovered reticulate histories for all species, including one potential case of hybrid speciation. Our results support the hypothesis that river dynamics promote speciation and reveal that many rainforest taxa are micro-endemic, unrecognized and thus threatened with imminent extinction. We propose that Amazonian hyper-diversity originates in part from fine-scale barrier displacement processes –including river dynamics– which allow small populations to differentiate and disperse into secondary contact.

## Introduction

The lowland rainforests of the Amazon River basin are home to one of the most diverse ecosystems on Earth, harboring more than 10% of all named species concentrated into an area that represents only about 0.5% of the Earths’ land surface area. Major hypotheses regarding the origins and assembly of this biota focus on the extreme heterogeneity of Amazonian environments, including the dendritic architecture of river drainage networks, the perennial role of river capture dynamics in fragmenting and merging riverine ecosystems through time and space, and the great antiquity of these systems dating back tens of millions of years (*1, 2*). The Riverine Barrier Hypothesis (RBH) posits that rivers can serve as barriers to dispersal and gene flow, fragmenting populations and causing diversification in many terrestrial organisms (*3*). Indeed, many rainforest assemblages of birds, primates, fishes, and other organisms exhibit high turnover in species composition on either side of large lowland Amazonian rivers (>1000m width at low water; Strahler Stream orders > 5), which dissect the whole region into broad interfluvial areas of endemism (*4*–*6*).

However, a burgeoning body of data indicates a more complex role for riverine barriers in generating patterns of Amazonian species richness. For example, comparative phylogeographic studies have shown that community-wide divergences across putative river barriers can be asynchronous (*7*), suggesting that divergence is instead driven by some combination of factors that include infrequent dispersal across pre-existing barriers (*7*), environmentally-mediated dispersal (*8, 9*), and ecologically-mediated divergence (*10, 11*). Although the debate over barrier-causality has often been framed in the context of dispersal versus vicariance (*12*), evaluating the RBH is complicated by the fact that barriers often change their location and permeability through geological time and across biogeographic space (*13*–*17*). Contemporary geographic features can therefore be poor indicators of past landscapes, a complication that has important consequences if trying to infer historical processes (*18*). For example, the river drainage network of lowland (below 250-300m elevation) Amazonia is now understood to be highly dynamic, with mega-river capture events of more than 10,000 sq. km occurring on timescales of tens to hundreds of thousands of years (*19, 20*). This dynamic landscape may be the primary mechanism generating patterns of aquatic biodiversity (*1*), but given such instability, some observers question whether Amazonian river courses actually persist long enough to act as effective barriers to gene flow for terrestrial organisms (*13*).

Despite much progress in understanding the processes mediating Amazonian diversification over time, a detailed picture of how biodiversity arises in this species-rich region is lacking. Most groups of Amazonian organisms lack genetic data with sufficient spatial resolution to discern the relationships between macroevolutionary processes (e.g. speciation and extinction) at biogeographic scales and microevolutionary processes (e.g. selection, migration, and drift) operating at populational scales (*21, 22*). To that end, many studies have identified patterns of differentiation that occur within the large interfluves (areas between major rivers), rather than across the major tributaries, including for birds (*23*–*26*), primates (*27, 28*), squamates (*29*), and butterflies (*30*). These patterns of differentiation have sometimes been attributed to isolation by smaller rivers (100–1000m width at low water)(*23, 26*), but other environmental factors that might affect population structure have rarely been tested. Thus, if biodiversity arises at micro-geographic scales, which fine-scale environmental features, if any, drive population differentiation, and do these features promote diversification via pure isolation or isolation-with-gene-flow?

The River Capture Hypothesis (RCH) posits that river network rearrangements (i.e. changes in riverine connections due to river captures and avulsions) that occur at appropriate spatial and temporal scales can drive diversification by increasing opportunities for speciation and dispersal, and thereby reducing the probability of extinction (*1*). To test this hypothesis, the southern Amazon Basin offers a unique test case. Southern Amazonia is characterized by a marked topographical gradient, in which sediment-rich rivers originating in the Andes in the west meet upland (above the fall line ∼250-300m elevation; Figure 1) rivers of the Brazilian Shield draining clearwater rivers from the southeast. These alternative geological settings, with rivers draining lowland-sedimentary and upland-incising basins, tend to be more and less dynamic, respectively (*31*). Sediment-rich lowland rivers frequently rearrange and continuously change via tributary captures and avulsions (*32*). Sediment-poor shield rivers rearrange less frequently and are characterized by climatically or tectonically driven headwater captures and recaptures (*19*). These two topographical regimes meet within the Madeira-Tapajós interfluve, a region that has been a hotspot of Plio-Pleistocene diversification (*8*). Importantly, sediment-poor rivers in this region may rearrange at intermediate rates; their upstream portions flow over the Brazilian Shield where they are relatively stable, but downstream portions flow into the lowlands, where they become more dynamic (*19*). The Madeira-Tapajós also marks an ecotone in forest structure driven by varying climatic regimes, wherein humid rainforests are more densely vegetated in western Amazonia but transition into more open forests to the south and east (*33, 34*). Given the many environmental gradients in Amazonia and its broad geographic extent, an important question is how river dynamics and environmental gradients influence demographic (e.g., gene flow) and phylogenetic (e.g., divergence and reticulation) histories in co-distributed Amazonian organisms.

**Figure 1:**
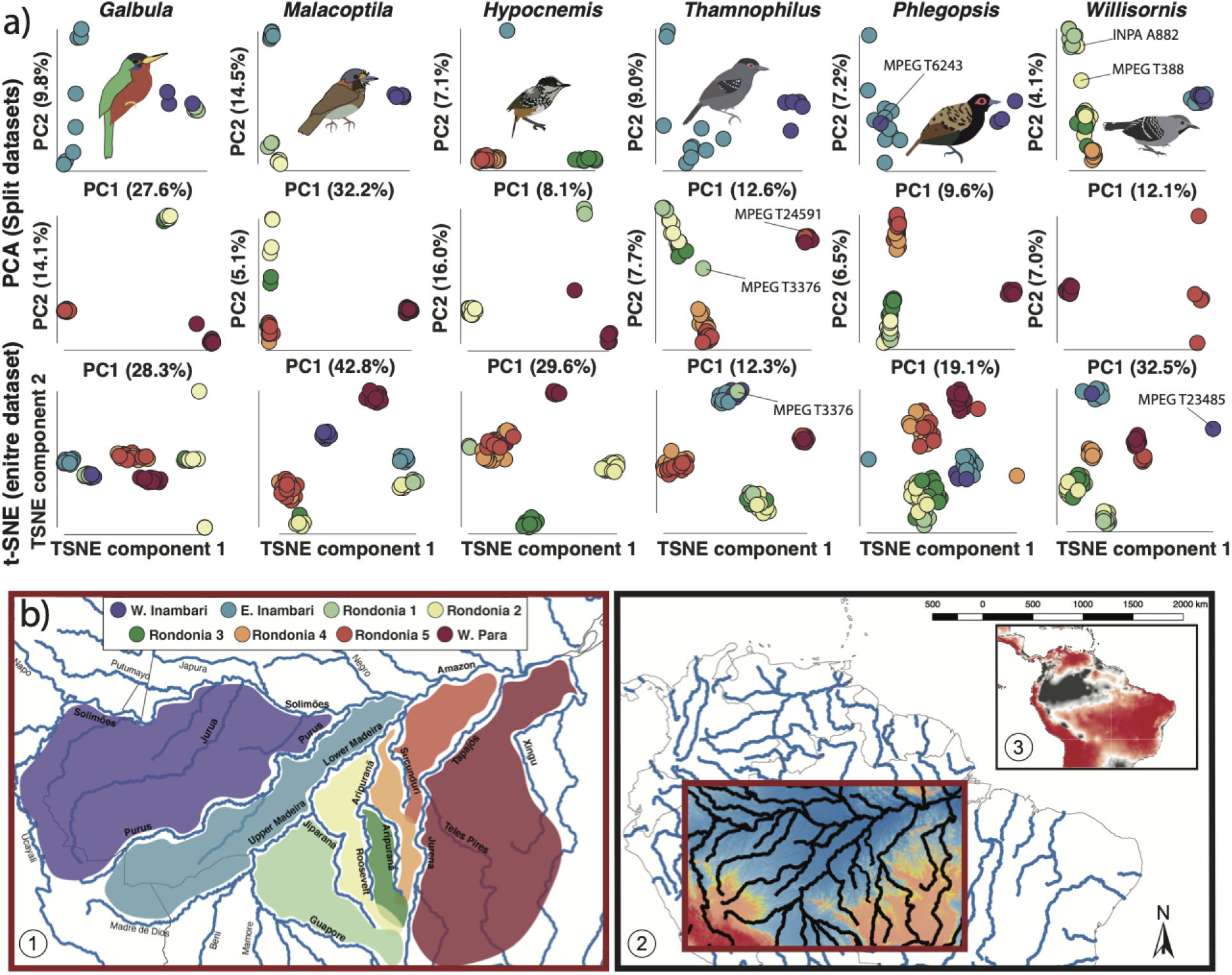
Population genomic variation in six co-distributed Amazonian bird species groups. (a) The top two rows show results from principal component analysis (PCA) where populations within each species-group were split into major divisions recovered in the species tree analysis (see Figure 2). The bottom row of plots shows example replicates from the t-distributed stochastic neighbor embedding (t-SNE) analysis for all individuals within each species. (b) Circles on the plots are colored based on major interfluvial regions in Map 1 (bottom left). Map 2 (bottom right) shows the topography of the study region within the dark red box, where yellow contours represent the 250–300m elevational zone demarcating dynamic lowland (<250m; shaded in blue) from relatively stable upland (>300m; shaded in red) basins. Map 3 (inset) shows precipitation during the driest annual quarter (*79*), with high precipitation in gray, low precipitation in red, and a strong cline across the middle to lower reaches of the rivers draining the Brazilian Shield (plotted using QGIS).

**Figure 2:**
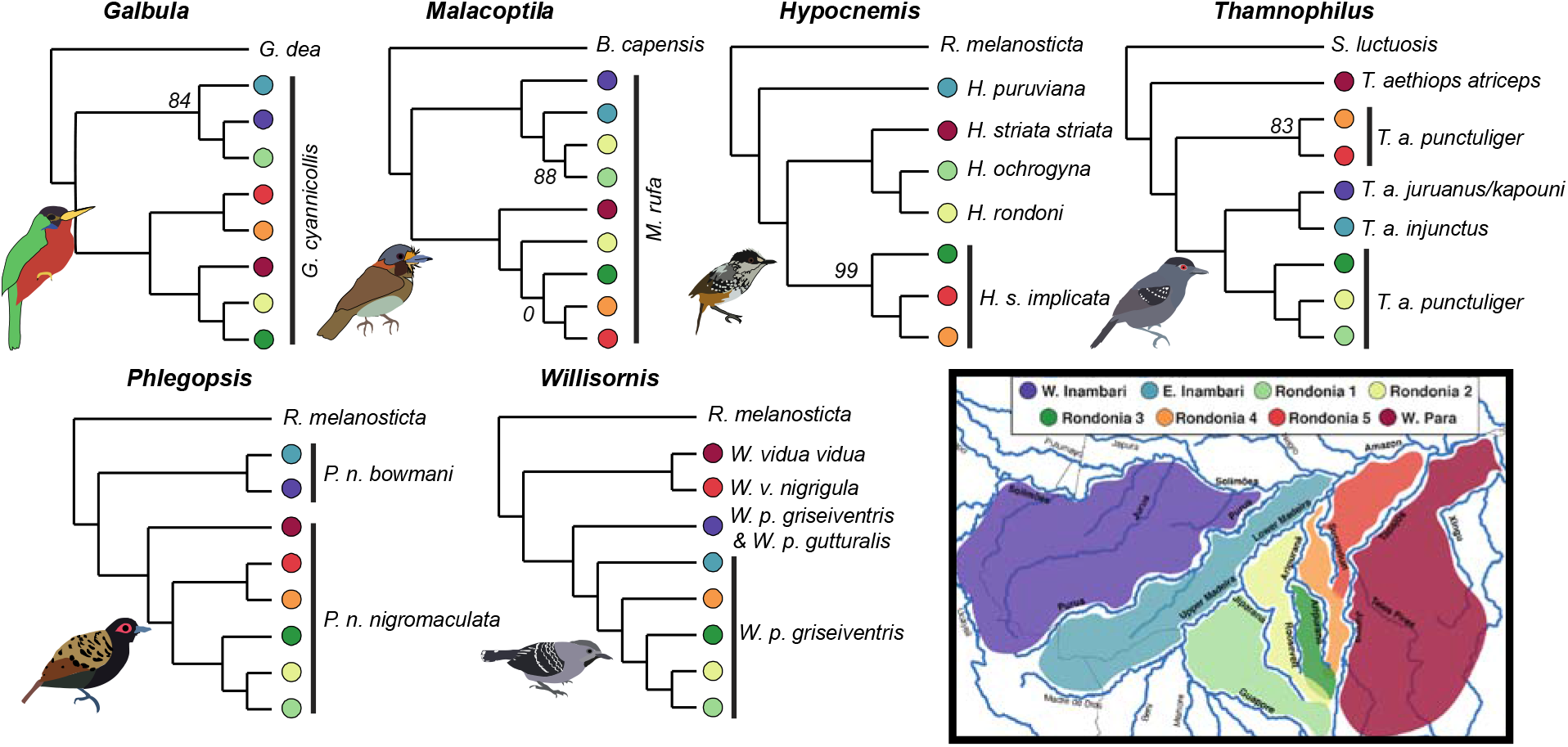
Summary of phylogenomic results for six Amazonian bird species groups examined in this study. (a) Results of species tree analysis in ASTRAL(*41*) showing the historical relationships among *a priori* defined populations in each species inferred from thousands of gene-trees. All nodes are recovered with 100% bootstrap support except where noted.

In this study, we sampled bird populations across the lowlands of southern Amazonia and sequenced hundreds of thousands of genetic markers to generate a dataset of 371 individuals sampled across six independently evolving species-groups. These groups vary in the amount of phenotypic divergence from monotypic species with little variation (*Galbula cyannicollis* and *Malacoptila rufa*) to polytypic species (*Thamnophilus aethiops* and *Phlegopsis nigromaculata*) or species-complexes (e.g., *Hypocnemis rondoni, H. striata, H. ochrogyna*, and *H. peruviana*., or *Willisornis poecilinotus* and *W. vidua*) with considerable phenotypic and behavioral variation. Despite their differences, all these taxa share relatively limited dispersal capacities and are tied to upland forests, making them good subjects for studying the effects of river dynamics. Our objectives are to (i) characterize genome-wide levels of genetic diversity in co-distributed avian species-groups, (ii) test the drivers of population genomic divergence in each species-group, (iii) quantify each species’ history of differentiation and gene flow, and (iv) test the hypothesis that river rearrangements promote diversification and gene flow in Amazonian birds.

### Assumptions, hypotheses, and predictions

Our hypotheses are predicated on a growing body of evidence suggesting that lowland Amazonian river networks are more dynamic over relatively short time-scales of tens to hundreds of thousands of years than are upland (shield) reaches of the same river basins (*13*). Since river rearrangements represent both the genesis and elimination of barriers to dispersal and gene flow, events that occur at appropriate spatiotemporal scales may leave signatures in the diversification histories of taxa (*20*). Therefore, if upland rivers are barriers to terrestrial organisms, then rearrangements among these watercourses would have important populational and phylogenomic consequences. For example, frequent river rearrangements, such as those expected in lowland sedimentary basins, may hamper population differentiation by homogenizing allele frequencies among populations on either side of a river (*35*). On the other hand, rearrangements may promote diversification by facilitating secondary sympatry between populations that had previously differentiated in allopatry, including for example, cycles of isolation with infrequent introgression (i.e. reticulation)(*36*). Thus, if river dynamics inhibit diversification, current river courses should not be strong predictors of population genomic structure (*13*). Instead, other processes may be better predictors of genomic variation. For example, isolation-by-distance (IBD, in which genetic differentiation increases with increasing geographic distance) (*37*) and isolation-by-environment (IBE, in which genetic differentiation increases with increasing environmental disparity) (*38*) are deviations from panmixia that can mimic the population genetic predictions of isolation-by-barriers (i.e., allopatry). If riverine barriers *do* structure populations, then river-course instability could stimulate diversification by promoting episodic isolation and secondary contact. In these cases, populations are expected to be structured across riverine barriers, but also show evidence of varying degrees of gene flow or sympatry between differentiated populations.

## Results

Our analyses revealed fine-scale population structure indicating a strong effect of current river channels in structuring genetic diversity. We used restriction site-associated DNA sequencing (RADSeq) to sample genome-wide sequence data for six avian species-groups co-distributed across eight populations in the southern Amazonian lowlands, and recovered high quality sequences for all six species (Tables 1 and S1). We defined these populations *a priori* as (1) West Inambari: south of Solimões River and west of Purus River, (2) East Inambari: Purus-Madeira interfluve, (3) Rondonia 1: Jiparaná-Guaporé interfluve, (4) Rondonia 2: Jiparaná-Roosevelt interfluve, (5) Rondonia 3: Roosevelt-Aripuanã interfluve, (6) Rondonia 4: Aripuanã-Sucunduri interfluve, (7) Rondonia 5: Sucunduri-Tapajós interfluve, and (8) West Pará: Tapajós-Xingú interfluve (Figure 1c Map 1). These seven regions were selected because they represent major interfluves bordered by rivers proposed to act as barriers (*23*–*26*).

**Table 1:** Assembly statistics for all species-groups, including the number of individuals in the assembly (n), the number of loci in the assembly (total loci), the average number of loci per sample (mean loci), the range of loci per sample (range loci), and the species of the reference genome sampled.

| **Taxon** | **n** | **Total Loci** | **Mean Loci** | **Range loci** | **Reference Genome** | **Genome Citation** |
| --- | --- | --- | --- | --- | --- | --- |
| *Galbula cyanicollis* | 53 | 87,606 | 46,669.10 | 13585–60141 | *Galbula dea* | Koepfli et al. (2015) |
| *Malacoptila rufa* | 51 | 64,477 | 31,179.70 | 16035–42638 | *Bucco capensis* | Koepfli et al. (2015) |
| *Thamnophilus aethiops* | 64 | 119,998 | 47,008.50 | 4089–68972 | *Sakesphorus luktusosis* | Koepfli et al. (2015) |
| *Hypocnemis sp.* | 61 | 110,035 | 52,986.10 | 9913–78143 | *Rhegmatorhina melanosticta* | Coelho, Musher, & Cracraft, 2019 |
| *Phlegopsis nigromaculata* | 77 | 122,081 | 64,081.80 | 19603–83765 | *Rhegmatorhina melanosticta* | Coelho, Musher, & Cracraft, 2019 |
| *Willisornis sp.* | 83 | 143,810 | 64,410.70 | 14266–81671 | *Rhegmatorhina melanosticta* | Coelho, Musher, & Cracraft, 2019 |

After characterizing genomic diversity, we recovered population structure associated with river barriers in all six species (Figure 1a). The first two axes of a principal components analysis (PCA) varied in their explanatory power across species groups, ranging from explaining 15.2% to 47.9% of total genetic variation in each dataset. To elucidate potential finer-scale patterns of differentiation in these species, we also utilized t-distributed stochastic neighbor embedding (t-SNE), a machine-learning dimensionality-reduction algorithm that is capable of detecting subtle characteristics of datasets (*39*). Although both PCA and t-SNE revealed similar results, t-SNE recovered additional structure in *Willisornis*, not detectable in the first two PC axes (Figure 1a). This included the separation of one sample, T23485, which represents a distinct subspecies from within the West Inambari region (*W. p. gutturalis*). An independent method for assigning individuals to ancestral populations, STRUCTURE v2.3.4 (*40*) showed less geographic structuring overall but recapitulated the general pattern of rivers as barriers (Figure S1). In all analyses, however, a handful of individual samples clustered outside of their geographic populations, a pattern possibly indicative of recent dispersal events.

Although all species were finely structured across space, we recovered unique spatial histories for each species-group (Figure 2). Species tree analyses (*41*) recovered robust phylogenetic topologies but differing area relationships among taxa. Bootstrap support for the relationships in these trees was generally high (>99), but the placement of the Rondonia 3 population in *Malacoptila* was unsupported. As the barrier associated with the deepest split in each taxon varied from large rivers such as the Madeira (*Hypocnemis* and *Phlegopsis*) and Tapajós (*Thamnophilus*) to smaller rivers such as the Aripuanã/Roosevelt (*Malacoptila*) or Sucunduri (*Willisornis*), these results are consistent with a scenario of isolation-by-barriers, supporting the notion that rivers of various sizes are historically important for population divergence across the central Amazon.

Interestingly, in *Malacoptila* we identified two apparently reproductively isolated, secondarily sympatric populations in the Rondonia 2 area (Figure 1a–b). Remarkably, these two overlapping populations were also syntopic, as they included two individuals, one from each population, collected at the same locality and date (MPEG T474 and MPEG T476). They are distantly related within this species-group (Figure 2) indicating secondary contact between divergent taxa despite strong isolation-by-barrier, a key prediction of the RCH.

### Spatially explicit models of population connectivity and genetic diversity

We found that populations in dynamic regions west of the Madeira river as well as near to the mouth of the Tapajós river likely experience higher rates of secondary contact (*42*). Whereas areas of low effective migration (hereafter, gene flow) relative to expectations under pure IBD overlapped with riverine barriers, in contrast, gene flow tended to be higher west of the Madeira where rivers are most dynamic (Figure 3). In three groups (*Galbula, Phlegopsis*, and *Willisornis*), we recovered evidence of higher genetic dissimilarity than expected under IBD west of the Madeira (Figures 4 and S3) despite high effective migration (Figures 3 and S2), a pattern indicative of secondary contact. We also found evidence of high migration coupled with high dissimilarity west of the mouth of the Tapajós river and north of the Aripuanã river for *Hypocnemis* and *Thamnophilus*. These results show that populations in these dynamic regions were more likely to experience secondary contact, a result consistent with predictions of the RCH.

**Figure 3:**
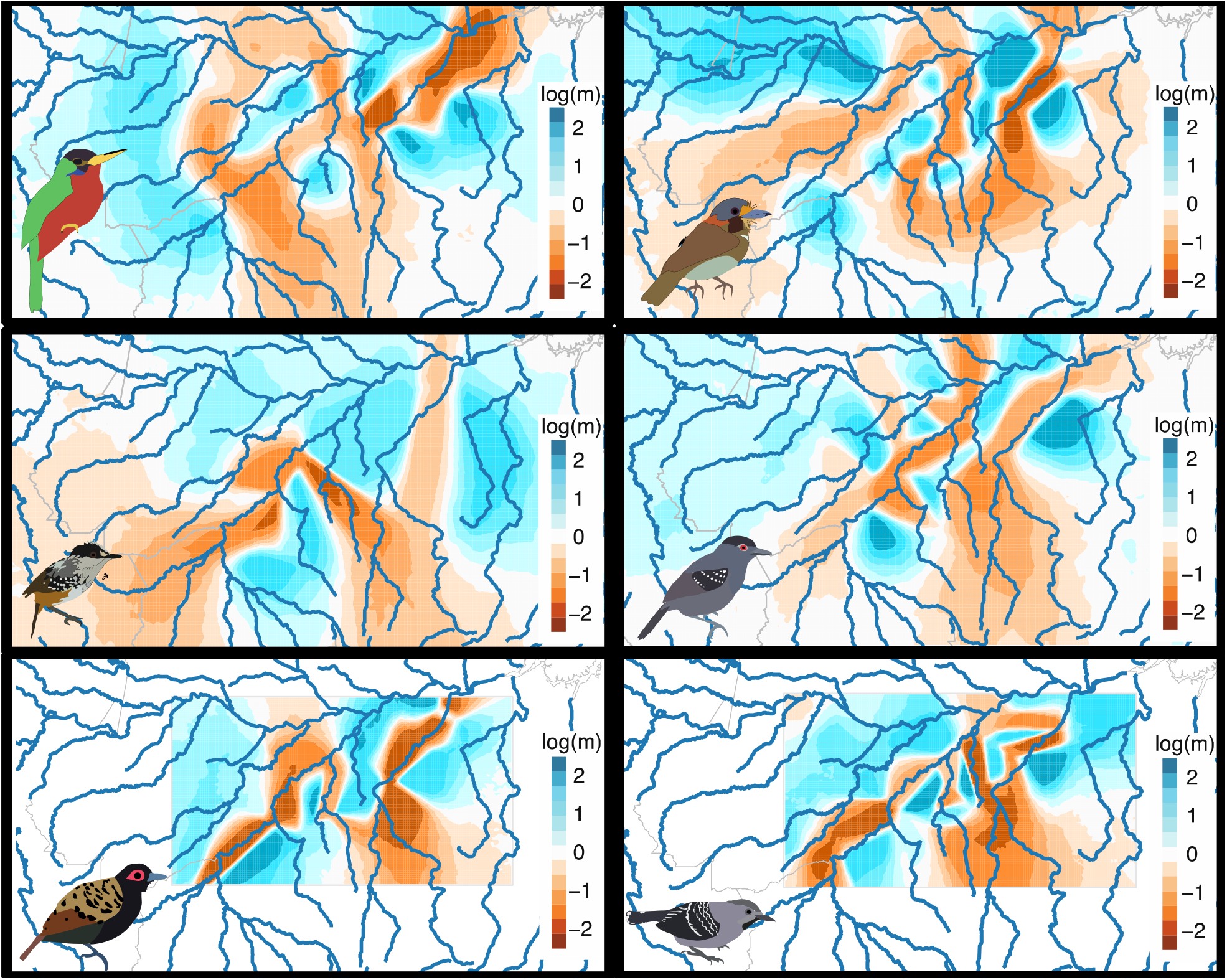
Effective migration results estimated in EEMS for (left to right, top to bottom) *G. cyannicolis, M. rufa, Hypocnemis spp*., *T. aethiops, P. nigromaculata*, and *Willisornis spp*. Effective migration rate (m) is shown on a log10 scale relative to the expected value under an IBD model across the sampled range. Darker blues correspond to higher effective migration rate, whereas darker oranges correspond to lower rates.

**Figure 4:**
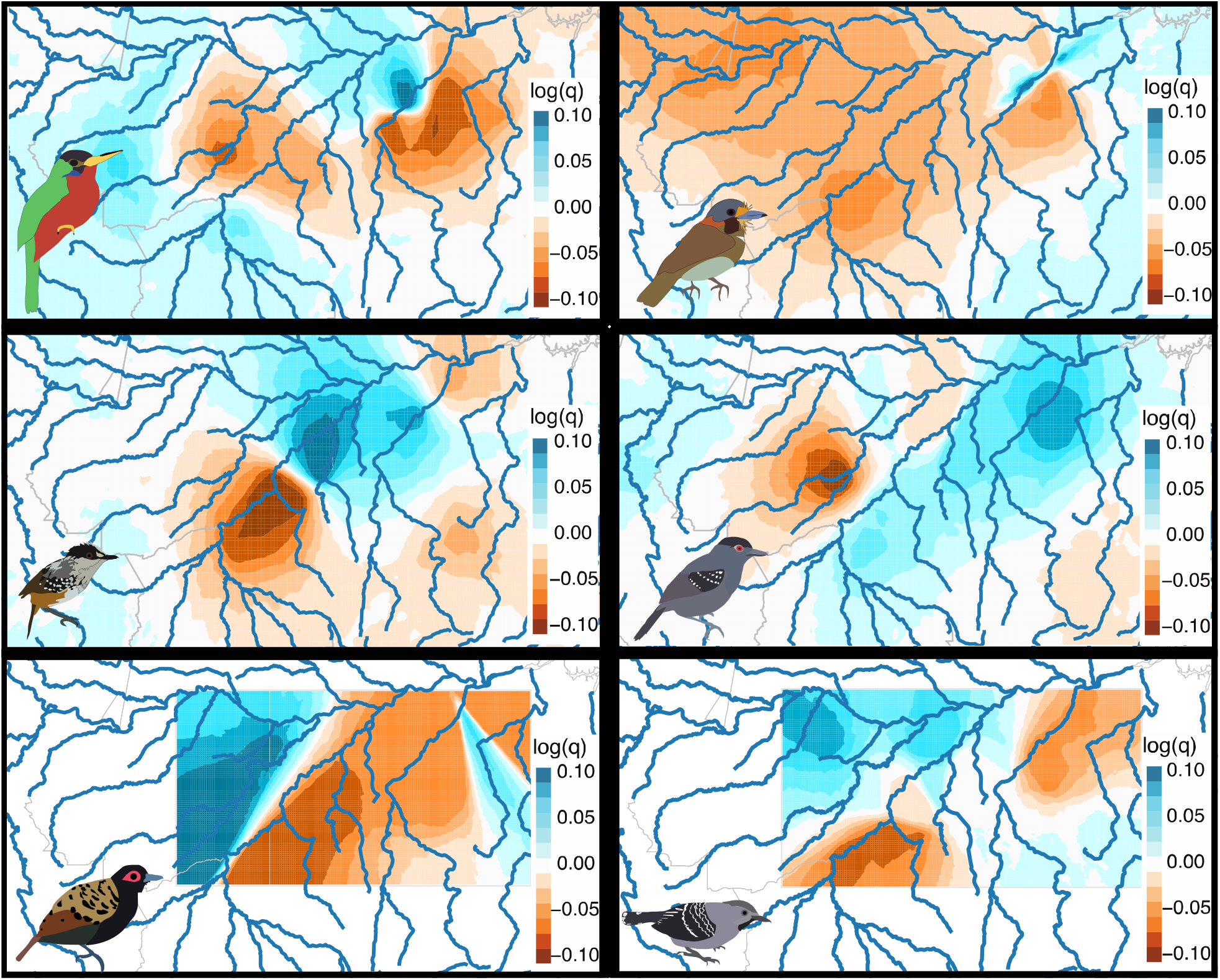
Effective diversity results estimated in EEMS for (left to right, top to bottom) *G. cyannicolis, M. rufa, Hypocnemis spp*., *T. aethiops, P. nigromaculata*, and *Willisornis spp*. Effective diversity (q) is shown on a log10 scale relative to the expected value under an IBD model across the sampled range. Darker blues correspond to higher genetic dissimilarity, whereas darker oranges correspond to lower dissimilarity.

### The predictors of genomic divergence

A multivariate model incorporating geographic dispersal distance (least cost-path), environmental disparity, and rivers rejected hypotheses of IBD and IBE as the primary drivers of divergences. Specifically, we found that river-barriers were significant predictors of genomic divergence in all groups irrespective of multicollinearity among predictor variables (Figure S4; Table 2) (*43*). The multivariate models strongly predicted genomic differentiation in each species-group, accounting for between 21% and 58% of the total variance in genomic divergence. Although the unique effects of geographic distance were relatively important for some species (explaining up to 20% of the R^2^), rivers were the most important predictors of genomic divergence in all six species-groups, explaining 27%–69% of the total variance explained by each species’ model (Table 2). Environmental disparity, though, tended to explain very little of each model (0.1%-8.6%). Thus, models explicitly testing the effects of IBD and IBE support the notion that rivers drive genomic divergence, a key prediction of the RBH and RCH.

**Table 2:** Results of the multivariate regression and commonality analysis, including beta-weights (β), p-values (*p*), and the coefficients for unique, common, and total contributions of each predictor. Per cent contributions to the model are shown in parentheses.

| Species | Model R-squared | Predictor | $\beta$ | structure coefficient | $p$ | Unique | Common | Total |
| --- | --- | --- | --- | --- | --- | --- | --- | --- |
| *Galbula* | 0.58 | DIST | 0.19 | 0.56 | <0.01 | 0.023 (3.9%) | 0.161 (27.8%) | 0.184 (31.7%) |
|  |  | ENV | 0.09 | 0.52 | <0.01 | 0.005 (0.8%) | 0.151 (26.0%) | 0.155 (26.8%) |
|  |  | RIV | 0.64 | 0.95 | <0.01 | 0.365 (63.1%) | 0.156 (26.9%) | 0.52 (90.0%) |
| *Malacoptila* | 0.52 | DIST | 0.14 | 0.54 | <0.01 | 0.123 (2.4%) | 0.138 (26.5%) | 0.151 (30.2%) |
|  |  | ENV | 0.03 | 0.44 | 0.25 | 0.001 (0.1%) | 0.101 (19.4%) | 0.102 (19.6%) |
|  |  | RIV | 0.65 | 0.98 | <0.01 | 0.358 (68.8%) | 0.14 (26.9%) | 0.498 (95.8%) |
| *Hypocnemis* | 0.28 | DIST | 0.27 | 0.55 | <0.01 | 0.033 (11.8%) | 0.050 (18.5%) | 0.083 (29.6%) |
|  |  | ENV | -0.22 | 0.20 | <0.01 | 0.024 (8.6%) | -0.013 (-4.6%) | 0.011 (3.9%) |
|  |  | RIV | 0.46 | 0.94 | <0.01 | 0.175 (38.0%) | 0.068 (24.3%) | 0.243 (86.8%) |
| *Thamophilus* | 0.21 | DIST | 0.20 | 0.64 | <0.01 | 0.026 (12.5%) | 0.061 (28.6%) | 0.087 (41.4%) |
|  |  | ENV | -0.04 | 0.39 | 0.16 | 0.001 (0.4%) | 0.032 (15.2%) | 0.033 (15.6%) |
|  |  | RIV | 0.37 | 0.92 | <0.01 | 0.123 (58.5%) | 0.056 (26.7%) | 0.179 (85.2%) |
| *Phlegopsis* | 0.34 | DIST | 0.27 | 0.69 | <0.01 | 0.047 (13.7%) | 0.116 (34.9%) | 0.1621 (48.6%) |
|  |  | ENV | -0.09 | 0.36 | <0.01 | 0.005 (1.5%) | 0.0384 (11.4%) | 0.0434 (12.8%) |
|  |  | RIV | 0.92 | 0.93 | <0.01 | 0.175 (51.8%) | 0.114 (33.7%) | 0.290 (85.7%) |
| *Willisornis* | 0.49 | DIST | 0.42 | 0.85 | <0.01 | 0.098 (20.0%) | 0.25 (51.0%) | 0.35 (71.0%) |
|  |  | ENV | 0.05 | 0.63 | <0.01 | 0.002 (0.0%) | 0.19 (38.8%) | 0.19 (38.8%) |
|  |  | RIV | 0.39 | 0.79 | <0.01 | 0.13 (27.5%) | 0.17 (34.7%) | 0.31 (63.3%) |

### Tests of introgression

Phylogenomic network analyses (*44*) recovered histories of introgression during diversification of these taxa despite clear spatial structuring around rivers, consistent with the RCH prediction that divergent populations experienced secondary contact (Figure 5a). We specifically recovered two reticulation events –the maximum allowed in the analysis– in all species-complexes but one (*Willisornis*). In most species, the most credible network topology was highly supported (PP=1.0), but the best models for *Malacoptila* and *Thamnophilus* were less supported. Unlike in other species, the dominant network topologies for *Hypocnemis* and *Thamnophilus* groups were inconsistent with the results of the species tree analysis, which suggests that ancestral or ongoing gene flow among differentiating populations has obscured species tree inference for these taxa. Estimates of interpopulation migration (gene flow) using a maximum likelihood population graph (*45*) were mostly consistent with the network modeling results, though with some notable differences (Figure 5b). The best-fitting model included at least one migration edge in each species but with diminishing gains in likelihood after that (Figure 5c). Thus, many adjacent populations separated by rivers likely experienced significant post-isolation gene flow.

**Figure 5:**
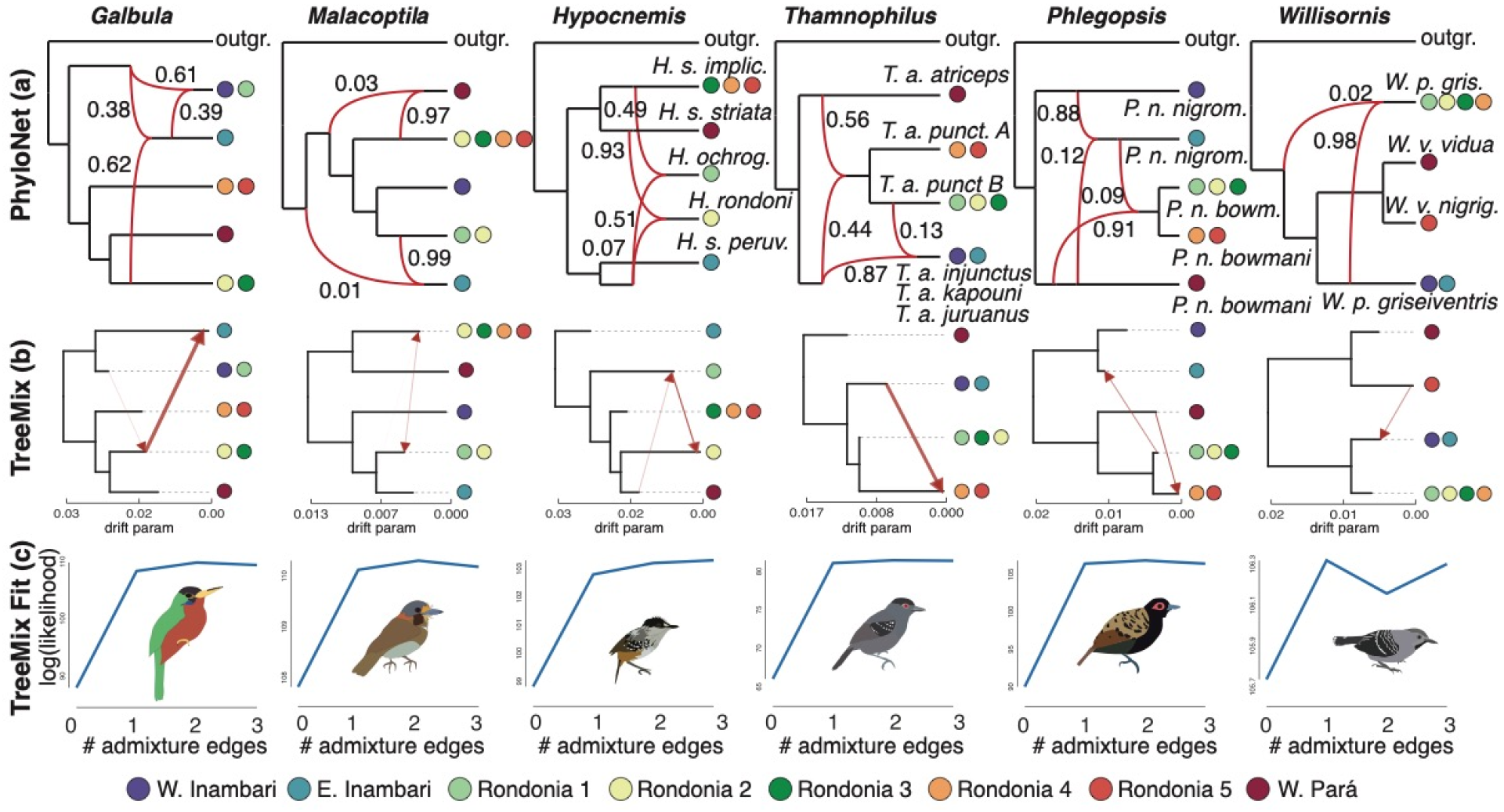
Summary of phylogenomic network results for six Amazonian bird species groups examined in this study. (a) Results of Bayesian Network analysis in PhyloNet (*44*) showing the most credible topology for each species group. Reticulate branches are shown in red and labeled with inheritance probabilities. Tips are labeled with taxonomic designations for all polytypic species-groups. (b) Results from TreeMix showing the inferred topology among populations (black branches) and inferred admixture edges (red arrows). The thickness of each migration edge is proportional to its inferred magnitude. (c) Results of the model-fitting exercise in TreeMix. The y-axes show the average log(likelihood) among all TreeMix replicates that varied in missing data thresholds for each value of m (the number of admixture edges). Circles at the tips of all networks are colored based on *a priori* defined populations (Figure 1 Map 1). Because PhyloNet and TreeMix populations were assigned using the results from STRUCTURE(*45*), multiple *a priori* populations may be sampled in a given tip.

Our analyses also revealed evidence of lineage fusions, implying portions of divergent populations have merged in the past. Within multiple species, at least one daughter node was recovered showing nearly equivalent inheritance probabilities from each of two parent nodes (*Galbula, Hypocnemis*, and *Thamnophilus*), where inheritance probability represents the proportion of sampled genes inherited through gene flow (Figure 5a). For example, the genome of *H. ochrogyna*, a phenotypically and behaviorally distinct taxon, resulted from nearly equivalent inheritance probabilities of parent nodes of different taxa, *H. striata* (p=0.49) and *H. peruviana* (p=0.51), a scenario consistent with hybrid speciation (*46*). Similarly, *T. a. punctuliger* has nearly equal shared ancestries with taxa in East and West Inambari (*T. a. kapouni, T. a. juruanus*, and *T. a. injunctus*) and Pará (*T. a. atriceps*). Within *G. cyanicollis*, multiple reticulate (non-bifurcating) nodes with high inheritance probabilities were associated with populations isolated by the Madeira and Purus, rivers known to have rearranged historically (*13*). Because incomplete lineage sorting is theoretically accounted for in the Multi-species Network Coalescent model, these shared ancestries are presumably driven by introgression. Thus, we reveal an additional distinct outcome of certain river network rearrangements, lineage fusion that results in genetically distinct lineages from the introgressing populations.

### Demographic modeling

Demographic modeling (*47*) revealed that all divergences likely occurred within the past 2my and that all extant populations were <1my old (Figure S5). However, divergences across the same river were asynchronous. Splits across the Tapajós formed two general groups; one at about 700kya to 1mya, and another at about 300kya (Figure S5b). Splits across the Madeira also roughly formed two groups, with most divergences being <1mya (Figure S5c). Across the Aripuanã-Roosevelt basin, divergences were mostly between 250kya and 500kya, except for *Phlegopsis*, which occurred about 70kya (Figure S5d). Although only three of the groups were differentiated across the Purus, two of these (*Galbula* and *Malacoptila*) diverged about 300kya (Figure S5e).

## Discussion

We demonstrated that the spatial distributions of genomically-characterized lineages within six Amazonian bird clades (i.e, species-groups) that inhabit *terra-firme* (non-floodplain) rainforests in southern Amazonia are delimited by the current position of many rivers, each with distinct hydrological and geomorphological profiles. Because existing river courses are the primary predictors of genomic divergence patterns in these species groups (*7*), our findings support the hypothesis that river dynamics promote diversification in southern Amazonia. Further, these results suggest that a significant portion of Amazonian biodiversity may be a product of historical barrier instability and dynamism across multiple scales.

Herein, we reveal that establishing secondary sympatry has had several distinct biological outcomes, including coexistence (*Malacoptila*), introgression (most groups), and possibly hybrid speciation (*Hypocnemis*)(*48*). For example, *Malacoptila rufa* experienced limited gene flow among differentiated populations, but secondarily sympatric populations in the Rondonia 2 area remained differentiated, indicating that these two populations are reproductively isolated despite any known phenotypic divergence (*23*). In other species, populations remained differentiated across rivers despite significant introgression from non-sister lineages (e.g., *Galbula, Thamnophilus*, and *Phlegopsis*). River dynamics may have even driven homoploid hybrid speciation, wherein *H. ochrogyna* –a widely recognized biological species (*25*)– resulted from an ancestral fusion between populations of *H. peruviana* and *H. striata*. Taken together, these results expose the necessity of accounting for gene flow when estimating evolutionary relationships in Amazonian taxa (*49*).

Homoploid hybrid speciation is thought to be rare in vertebrates, but recent studies have highlighted an increasing number of examples (*46, 50*). Indeed, lineage fusions, wherein diverging lineages –or portions of those lineages– merge to become geographically isolated but admixed evolutionary entities, seem to be more common than previously realized (*51, 52*). These fusions have been proposed to occur due to a number of biogeographic mechanisms that include river rearrangements in Amazonia (*46*). The examples identified herein add to this growing body of evidence that the disappearance of physical barriers to gene flow can facilitate the merging of divergent populations, that may then emerge as novel admixed taxa (*53*).

Importantly, the robustness of our results is independent of the taxonomic approach adopted. If adopting a phylogenetic species concept, for example, continuous river dynamism may repeatedly isolate novel sets of lineages that become differentiated, irreducible clusters of individuals (e.g., Figure 1)(*54*). Under a biological species concept (*55*), cycles of isolation and of secondary sympatry in this context can lead to reproductive isolation, either through the buildup of postzygotic barriers (*56*) or possibly adaptive introgression (*36*). To be sure, *H. ochrogyna* and *H. rondoni* are phenotypically and vocally distinct, and are reproductively isolated, yet admixed species (Figure 5a–b)(*25*). *Hypocnemis ochrogyna* is known to be isolated from *H. striata* via postzygotic barriers, and a similar process may drive isolation between other sets of neighboring taxa in this genus (*56*) and others as well, including *Willsornis* (*57*). We therefore suggest that changes to the positions of rivers can result in cohesive and isolated species boundaries despite substantial ancestral admixture (*58*).

### Bridging diversification models of aquatic and terrestrial Amazonian organisms

Spatiotemporal patterns of biodiversity are shaped by geophysically- and climatically-generated cycles of fission and fusion among populations, species, and biogeographic areas (*59*). These episodic cycles play out across multiple scales and with varying results across the globe as ecosystems expand and contract, barriers to dispersal wax and wane, and as distributions change over time (*15, 60, 61*). At shallow evolutionary timescales, and as exemplified herein, these fission-fusion cycles have the power to generate diversity by driving differentiation via isolation, colonization, and even genomic and phenomic recombination (i.e., adaptive introgression) (*36, 51*).

Our study exemplifies a possible geomorphological mechanism –river drainage evolution and rearrangement– through which these fission-fusion cycles may play out for Amazonian organisms that rarely cross rivers and their associated floodplains (*62*). Like other fission-fusion cycles, rates of river rearrangement are ultimately controlled by tectonic, climatic, and erosional processes that have myriad effects on species diversification (*14, 19, 31*). For example, the process of barrier displacement (i.e., the movement of barriers to gene flow over time)(*60*), is thought to have had a profound effect on freshwater fish diversity, which reaches its global zenith in Amazonia (*6*). Fish alpha diversity (local diversity) is highest in the lowland sedimentary basins, where the low topography facilitates frequent connections and disconnections among watersheds driven primarily by erosional forces. In contrast, fish beta diversity (endemism) is higher in the upland shields where rearrangements occur less frequently (*5*). To summarize, more frequent river capture events facilitate higher rates of both isolation and dispersal (i.e., secondary sympatry) among adjacent watersheds, thereby promoting low spatial and high temporal fish species turnover in lowland western Amazonia, but high spatial and low temporal turnover in the upland portions of eastern Amazonia (*1*).

This core-periphery pattern is mirrored by broad patterns of many terrestrial organisms in two ways. First, phylogenetically under-dispersed communities of birds inhabiting the tectonically unstable western basin indicate rapid *in situ* diversification and the repeated establishment of secondary sympatry, while phylogenetically over-dispersed communities of the more stable Brazilian Shield suggest slower diversification with higher local endemism (*12, 31*). Second, rate estimates of phylogeographic lineage splitting for Amazonian birds is higher in the western lowlands than eastern shields. Rates of lineage loss, however, are overall higher in the uplands (*21*). Our results show that this pattern holds true for certain taxa at microevolutionary scales as well; effective diversity is higher across western Amazonia for three out of the six taxa despite higher rates of migration indicating more rapid episodes of isolation and secondary contact at the epicenter of drainage rearrangement (Figures 3 and 4).

Biogeographic simulations reveal that barrier displacement processes have distinct predictions when compared with purely stochastic processes (*60*). For example, barrier displacement results in high species richness and younger diversity toward the epicenter of displacement activity (e.g., Amazonia), but lower species richness and deeper-branching (i.e., less diversified) lineages toward the periphery of such landscapes. Our data focus primarily on the population genomic patterns within the Madeira-Tapajós of central Amazonia, and therefore do not directly test barrier displacement theory. However, in combination with previous work, our data lend support to this model in multiple ways. First, we show that the biogeographic processes acting on Amazonian birds are dominated by isolation-by-barrier, and not local adaptation to environmental differences, as environmental disparity explains little if any variation in genomic divergence for all sampled taxa (Figure S4; Table 2). Second, as mentioned above, a key prediction of barrier displacement theory is the buildup of older lineages at the continental periphery. This prediction is satisfied by many avian-species groups. For example, in *Thamnophilus* and *Malacoptila*, we confirm several young isolated lineages occur across the Amazon Basin, but previous work has documented deeper-branching, less diversified clades that occur at Amazonia’s periphery in the eastern Andean foothills and Atlantic Forests, respectively (*23, 24*). Likewise, in the clade of ant-following antbirds, of which *Willisornis* and *Phlegopsis* are a part, most diversity is restricted to lowland Amazonia, with the highest levels of sympatric species in the sedimentary basins, where riverine barrier displacement is most common (*63*). These sympatric taxa include close relatives within the same genus (e.g., *Phlegopsis and Rhegmatorhina*)(*63, 64*). The earliest branching clade in this group, *Phaenostictus mcleannani* plus *Pithys*, however, is distributed in northwestern South America, which suggests lower rates of net-diversification at the periphery (*63*).

Through this study, we catch a glimpse of how barrier displacement in the form of river rearrangement can facilitate isolation and secondary sympatry as populations of *M. rufa* in the Rondonia 2 area became syntopic and others show reticulation. For example, although no two species’ histories were phylogenetically congruent with respect to the details of geography, some general patterns emerged among species. All six species showed either phylogenetic (*Galbula, Malacoptila, Thamnophilus*, and *Willisornis* species trees) or reticulate (*Galbula, Malacoptila, Hypocnemis*, and *Thamnophilus* networks) affinities across the upper Madeira, a section of river thought to have been captured from a tributary of the Purus during the late Quaternary (Figures 2 & 5)(*13*). Taken at face value, our data suggest this capture event occurred roughly 100–800kya, consistent with or older than previous estimates (Figure S5c) (*13, 65*). Similarly, five of six species showed phylogenetic or reticulate affinities across the lower Tapajós, where avulsion shifted this downstream portion multiple times (*19*). Phylogenomic trees and networks show that the Para population is sister to northern populations of the Madeira-Tapajós for *Malacoptila* (Rondonia 2–5) and *Willisornis* (Rondonia 5), implying that these avulsions at the mouth of the Tapajós may have occurred at least twice: once at roughly 250kya for *Malacoptila*, and once around 600kya for *Willisornis* (Figure S5b). Moreover, in *Galbula* and *Hypocnemis*, the Para population is sister to populations at the center of the Madeira Tapajós (Rondonia 2 &3), where river capture is thought to have shifted drainage from Madeira tributaries to Tapajós basins (*5*). Overall, then, we provide evidence that drainage instability, rather than river formation *sensu stricto*, may drive diversification dynamics in many Neotropical organisms limited by rivers.

Therefore, the RCH as a general framework unifies biogeographic theory of both aquatic (primarily freshwater fish) and terrestrial (primarily birds and primates) Amazonian vertebrates, and suggests a common cause for species accumulation in these groups (*31*). First, macroevolutionary patterns of biodiversity seem to arise at microevolutionary scales (*21, 23, 26, 28*), and herein we demonstrated that even small rivers can be key drivers of microevolutionary diversity for many Amazonian birds (Figure S5; Table 2). Since lowland rivers are typically dynamic, we propose that, like many aquatic organisms, Amazonian terrestrial diversity originates from river dynamics at local to regional scales, which promote the early divergence of populations, forming spatially restricted and differentiated populations (i.e., taxa)(*20*). Ongoing river rearrangements may then promote dispersal, leading to secondary contact among new sets of populations, as well as new isolation and divergences. As these rivers shift course via river-capture and avulsion they directly affect the three fundamental parameters of diversification: dispersal (via the loss of barriers; i.e., geodispersal), speciation (via isolation and secondary contact), and extinction (via changes to population connectivity or spatial restriction) (*1*). Given Amazonia’s ancient history and probable low extinction rates (*21*), the repetition of this process through deep time has likely promoted the accumulation of a large portion of its vertebrate diversity. In this way, Amazonia is both a species-pump that continuously generates new diversity and also a ‘museum’ for old lineages that originated in or colonized the South American tropics long ago (*66*).

### The drivers of Amazonian endemism

Our results suggest that a long-recognized area of endemism (Rondonia *sensu* Cracraft (*4*)) may not represent a historically unified biogeographic unit for many taxa. Fernandes (*26*) argued that this biogeographic region instead contains multiple areas, each with a distinct history. Our data corroborate this notion revealing that endemism within Rondonia is more finely structured than currently recognized (*26*). In four of the six sampled species-groups, sets of populations within Rondonia were not reciprocally monophyletic. Rather, *Gabula, Malacoptila, Hypocnemis*, and *Thamnophilus* all included populations within Rondonia with sister relationships either west of the Madeira or east of the Tapajos, rather than to other Rondonian populations. Importantly, the patterns and processes we identified are likely not limited to the region’s avifauna; the distributions of primates in Rondonia are strikingly similar to, if not more spatially-restricted than, those identified herein (*67, 68*).

Consequently, areas of endemism in general may be individuated based on a misconstrued understanding of patterns of diversity. As we show, the histories of species in this study were characterized by gene flow between non-sister taxa (genomic reticulation) and phylogenetic incongruence with respect to the details of geography (biogeographic area reticulation)(*5*). Thus, Rondonia has been connected to and disconnected from other areas over evolutionary time, and as a result, the components of its biota are of different ages (*5, 7*). Although Rondonia contains a distinct contemporary biological community, it may be illusory as it is in part a construct of taxonomic artifact (unrecognized taxa at smaller spatial scales) and in part the result of multiple superimposed histories of isolation and expansion yielding quasi-congruent spatial patterns (*69*).

In general, then, areas of endemism may instead result because taxa originate at smaller scales and then expand their ranges as dynamic barriers to dispersal erode away. In regions with high rates of barrier displacement such as the western Amazon, the signal of fine-scale diversification and endemism may quickly erode away due to high temporal turnover (*12, 31*). In regions with somewhat slower rates of displacement, such as those within the Madeira-Tapajós, where shield and sedimentary rivers meet, the signal of local diversification may remain over longer periods, facilitating detection by studies such as ours (*12, 26*). Therefore, as Amazonian watersheds are reticulate biogeographic areas for aquatic organisms (*5*), so too might they be for endemism in birds. Fine-scale ephemeral processes may generate diversity, but this diversity eventually expands to reach barriers that are, though not static in the strict sense, less permeable at a given point in time (*31*). These dynamics may often, but not always, result in sister relationships across larger rivers over deeper timescales (*70*).

### Limitations and future directions

Although our findings support the RCH, the effects of rivers likely interact with other processes. For example, climate change can alter the strength of riverine barriers and can also drive local extinction dynamics, thereby mediating extinction-recolonization cycles (*8*). Such effects of climate on barrier effectiveness may be indistinguishable from river rearrangements in the strict sense based on the data we provide and could also explain some patterns found herein. Given that there are many young divergences across the Brazilian Shield (*8*), a broader sampling regime extending across lowland and upland basins in southern Amazonia may be necessary to disentangle the two processes with more certainty (*1*). Moreover, because we do not explicitly test pulse-migration versus continuous-migration models of gene flow, the process of happenstance dispersal across stable barriers may be difficult to distinguish from a barrier displacement model in practice. However, the explicit assumption of the framework presented herein is that the underlying lowland riverine landscape is not stable, which has also been confirmed, for instance, on the easternmost Amazonian part of the Brazilian Shield (*71, 72*). Within this conceptual framework, drainage network evolution is a simple solution to the question of how secondary contact occurs between differentiated taxa that are otherwise isolated by rivers.

### Concluding remarks

Amazonian biodiversity is unmatched by any other global ecosystem (*73*). Still, we demonstrate its species richness may be greatly underestimated even in well-studied groups such as birds (*2, 56*). This diversity, though, is non-randomly distributed across the basin, with community composition shifting across many rivers. Our results corroborate those of other studies that have found fine-scale endemism across the Madeira-Tapajós interfluve –a region threatened by rapid and ongoing deforestation (*26, 74*)– yet this endemism is generally unrecognized (*73*). The populations delimited by these rivers represent taxa, as they are differentiated, isolated evolutionary entities even if they are genomically admixed. They are relatively young (<500ky) and spatially restricted. That these taxa are micro-endemic conveys the severity of future biodiversity loss in southern Amazonia should deforestation continue at current levels (*75, 76*); that many of them are not yet named and formally described reveals how much fundamental biological information of the Amazonian avifauna is threatened with loss to imminent extinction.

## Materials and Methods

### Restriction site-associated DNA sequencing (RADSeq)

We extracted total DNA from vouchered fresh tissue samples using a DNeasy tissue extraction kit (Qiagen, Valencia, California), which recovered sufficiently high-quality DNA (>10kb fragments) for all samples. Library prep for restriction site-associated DNA sequencing (RADSeq) was then performed at the University of Wisconsin Biotechnology Center (UWBC, Madison, WI) using two enzymes (Pstl and Mspl) but barcoding only a single cut site (TGCA). 150 base-pair paired-end sequencing was then performed on an Illumina NovaSeq.

Raw Illumina reads were processed using iPyrad version 0.9 (*77*). We first demultiplexed the raw reads and trimmed low-quality base-pairs, and then aligned reads for each species to a reference genome. We specifically applied a minimum coverage for statistical base-calling of six, with minimum trimmed read length of 35 bp, a maximum of 5% uncalled bases and 5% heterozygous sites, and a mapping threshold of 0.9. These steps were repeated to generate variant call format files for each species group, which were used and filtered for downstream analyses.

### Characterization of genomic variation

We took multiple approaches to characterize genomic diversity across each species. First, we used iPyrad API analysis tools (*77*) to perform Principal Components Analysis (PCA) on each species. Because RADSeq datasets often contain a large proportion of missing data, we performed PCA after imputing missing haplotypes using k-means clustering implemented in the iPyrad API. This method allows imputation without *a priori* bias about population assignments. Specifically, we assumed k=8 for all species (except *Hypocnemis* because West Inambari was not sampled) and used iterative clustering to group individuals into populations. For iterative clustering, we first sampled SNPs present across 90% of individuals in the assembly, then clustered with the assumed K. This was repeated five times, allowing more missing data at each successive iteration until reaching a minimum of 75% coverage at each sampled SNP. To impute, we then randomly sampled haplotypes based on the frequency of alleles within each of the populations defined by k-means. Because each species was characterized by a large amount of total genomic divergence across PC space, we split each species’ dataset into two population subsets based on phylogenetic relatedness (i.e., clades; see ASTRAL analysis below), and ran PCA on each data subset to improve visualization. We also utilized t-SNE, which is explained in further detail below. As an independent method for assigning individuals to populations, we used STRUCTURE v2.3.4 (*45*) (for additional detail, see electronic supplementary methods).

### Spatially explicit models of population connectivity and diversity

To examine how the distribution of population genomic diversity within each species group varies across space, identify regions of high or low dispersal, and evaluate the contribution of IBD to spatial patterns of genetic variation, we used Estimated Effective Migration Surfaces (EEMS) (24). As input for EEMS, we first quantified Euclidian genetic distance matrices between individuals for each species complex using iPyrad API analysis tools (*77*). To generate polygons to constrain the spatial extent of the analysis, we drew a rectangle around all sampled localities for each species and distributed 1000 demes over each one. We then performed three runs of the Markov chain Monte Carlo (MCMC) for 2,000,000 iterations, thinning every 10,000 iterations, and excluded a burn-in of 1,000,000 iterations. The convergence of the MCMC was visually assessed by plotting the log posterior probabilities of each run and comparing results among runs for the same species. When results were converged among runs, a single run was chosen.

### Testing the predictors of genomic divergence

We tested for the effects of geographic distance, environmental disparity, and river barriers on genomic divergence in each species-group while accounting for spatial autocorrelation among these variables using commonality analysis. Because we were interested in fine-scale patterns of differentiation, we quantified pairwise genomic divergence among samples in each species using t-SNE (*39*) implemented in the iPyrad API analysis tools. t-SNE is a machine-learning dimensionality-reduction algorithm that decomposes high-dimensionality data into two components for more intuitive quantification and visualization, and is capable of detecting subtle characteristics of datasets (*39*). Here, t-SNE was used to quantify the total separation of samples across PC space (see above).

However, t-SNE results are sensitive to the perplexity (an approximate estimate of the number of neighboring points per cluster) and starting seed used. Although these values don’t typically influence the general patterns recovered by t-SNE, no two t-SNE runs that differ in seed and perplexity will produce identical results. To account for variance among t-SNE runs, we performed 10,000 t-SNE replicates, randomly choosing perplexity values (integers between three and eight) and starting seeds. We then used the average pairwise Euclidean distances between samples in t-SNE space as a measure of genomic divergence. While genetic distance or PCA can be used as measures of genomic divergence among samples, using t-SNE allows for evaluation of local-scale differentiation among individuals that may be too subtle to detect with other approaches, including PCA.

We measured geographic distance by taking the least cost-path distance between sample localities based on environmental niche models (ENMs; see Electronic Supplementary Methods). This method provides a metric of geographic distance that reflects the most likely dispersal paths based on habitat suitability on the landscape. The least cost path distances were calculated using the R package GDISTANCE (*78*). To measure environmental disparity, we extracted climatic data from 19 BioClim variables (*79*) for all sample localities, and used PCA to summarize the climatic conditions at each locality. We then calculated the Euclidean distance between all samples in this ten-dimensional climatic PC space.

Finally, to evaluate the effect of river barriers, we generated a pairwise matrix of individuals, where sample pairs were scored as either 0 (not separated by a river) or 1 (separated by a river). We specifically tested the seven river barriers of interest corresponding to the boundaries of the eight *a priori* defined populations (Purus, Madeira, Jiparaná, Roosevelt, Aripuanã, Sucunduri, and Tapajós Rivers). Although not all rivers should be considered equal, we discretized these barriers to conservatively test for their effects on genomic divergence in general. For our purposes we treat the Juruena and Teles-Pires rivers as synonymous with the Upper Tapajós, and refer to them this way throughout.

To evaluate the relative contributions of each variable in a multivariate linear regression while controlling for multicollinearity (i.e., spatial autocorrelation) among predictors, we used commonality analysis (*43*). Commonality analysis is a variance partitioning approach that determines the relative contributions of a set of independent variables (geographic distance, environmental disparity, and rivers) on a dependent variable (population genomic structure) while accounting for nonindependence among predictors. This helps to pinpoint the location and extent of multicollinearity among variables. To do this, a multivariate linear regression model is decomposed into its unique and common contributions. The unique contribution of a given predictor represents the proportion of the R^2^ (i.e., the proportion of total variance in genomic divergence explained by the multivariate linear regression model) that is explained solely by that predictor. Common contributions represent the proportions of the total R^2^ that are explained by that predictor in combination with other predictors. The total contribution of each predictor, the commonality coefficient, is thus quantified irrespective of any nonindependence among variables.

We used among-sample pairwise matrices of geographic distance, environmental disparity, and river barriers as predictor variables, and pairwise genomic divergence as the response variable. We also used a z-transformation (subtract the mean and divide by standard deviation) to standardize predictors (*43*). We calculated commonality coefficients and then computed 95% confidence intervals around those coefficients by bootstrapping with 1,000 replicates of a random selection of 90% of samples without replacement. We specifically adapted R scripts from Prunier et al. (*43*) and Seeholzer and Brumfield (*80*).

### Phylogenomic species trees

We estimated the historical relationships among *a priori* populations using ASTRAL 5.7.3, (*41*). ASTRAL estimates the species tree from a set of unrooted gene trees under the multi-species coalescent model, with an assumption of no gene flow among species. To assess robustness of the inferred population relationships, we also applied 100 bootstraps, subsampling gene trees only, using the *--gene-only* flag. Gene trees were generated for all loci greater than 400 bp in length. To reduce potential noise from gene tree estimation error, we then applied a filtering algorithm that identifies outlier loci (loci deviating from clock likeness expectations across the genome)(*61*). Species trees were rooted based on the reference genome sequences for each species-group (see ESM).

### Tests of introgression

To examine the evolutionary relationships within each species while relaxing the assumption of no gene flow, we additionally applied Bayesian phylogenomic network analysis using the MCMC_GT algorithm implemented in PhyloNet (*44*). This algorithm uses a reverse-jump Markov Chain Monte Carlo (rjMCMC) to sample the posterior distribution of phylogenetic networks under a Multi-Species Network Coalescent model (MSNC). The MSNC models genome evolution as a network (as opposed to bifurcating tree), accounting for both incomplete lineage sorting and inter-lineage gene flow (reticulation). We used the same set of gene trees filtered under above criteria to estimate the network for each species. We ran the rjMCMC for 5 × 10^7^ generations with a burnin of 5 × 10^6^, sampling every 10^5^ generations, using the pseudo-likelihood calculation, and allowing a maximum of two reticulate nodes per network (the rjMCMC will thus test networks with zero, one, and two reticulations only). The network topology with the highest posterior probability (PP) is chosen as the most credible network topology.

Finally, we used TreeMix 1.13 (*45*) to estimate the maximum likelihood population graph in the presence of gene flow by attaching migration edges (inferred shared ancestry due to genetic exchange) to a tree of bifurcating populations. Migration edges are assigned migration weights, which are related to the proportion of alleles in a population that are derived from migration. To examine the sensitivity of TreeMix output to missing data, we first ran TreeMix on subsamples of 50%, 60%, 70%, 80%, and 90% missing data with one migration edge (m=1). We then examined the model fit for varying values of m (m=0– 3), where m represents the number of migration edges in the model (zero represents a pure isolation model). To do so, we performed 250 TreeMix replicates, randomly choosing missing data thresholds between 50% and 95% and quantifying the likelihood of the model and averaging across all replicates. This was repeated for each value of m.

Because unstructured populations may experience high gene flow, for both PhyloNet and TreeMix we assigned individuals to populations based on their assignments in STRUCTURE. However, because STRUCTURE analyses resulted in underestimation of population structure for *Thamnophilus* and *Phlegopsis* when compared with PCA and previous work (*8, 24*), we split one of the population assignments in two based on PCA results with the goal of better understanding spatial patterns of isolation and reticulation in these groups (for additional detail, see electronic supplementary methods).

### Demographic Modeling

To estimate divergence times and effective population sizes in the presence of gene flow, we applied a Bayesian approach using Generalized Phylogenetic Coalescent Sampler (G-PhoCS) (*47*). G-PhoCS integrates full likelihood computation across all possible gametic phasings based on unphased loci and a predefined demographic model to estimate mutation-scaled effective population sizes (θ) and divergence times (τ) for all populations. The software also allows estimation of continuous gene flow between sets of current or ancestral populations. We constructed demographic models for each species-group based on the results from PhyloNet. Specifically, reticulate edges in PhyloNet with higher inheritance probabilities (>0.5) were treated as population divergences, whereas the edges with lower probabilities (<0.5), were treated as migration edges. We allowed for all possible migration edges based on both PhyloNet and TreeMix results. We applied priors on root τ (α = 3.0; β= 1,000) and θ (α = 5.0; β= 1000), and for each population and divergence (τ α = 1.0, β= 50,000; θ α = 5.0, β= 1,000). We then ran the MCMC for 5 × 10^5^ iterations after a burn-in of ten percent, sampling every ten iterations. To convert τ and θ outputs to absolute divergence times (T) and effective population sizes (Ne), respectively, we used the formulas Ne = θ/4μ and T = τG/μ, where μ is the mutation rate in substitutions per site per generation and G is the generation time in years (for additional detail, see electronic supplementary methods).

## Acknowledgements

We thank Tommaso Chiodo, Lauren Audi, Stephen Gaughran, Anthony Caragiulo, Jeff Groth, Paul Sweet, Romina Batista, Fátima Lima, Ivá Jean, and Leôncio Jean for advice and assistance at various stages of this work. We additionally thank Emily Grifith, Jon Merwin, Kamila Kuabara, Abigail Del Grosso, and Jason Weckstein for helpful comments on a previous version of this manuscript.

