## Supplemental Information for "River network rearrangements promote speciation in lowland Amazonian birds"

**Supplementary Information**

**Detailed Methods**

*Restriction site-associated DNA sequencing (RADSeq)*

We extracted DNA from vouchered fresh tissue samples using a DNeasy tissue extraction kit (Qiagen, Valencia, California), which recovered sufficiently high-quality DNA (>10kb fragments) for all samples. Library prep for restriction site-associated DNA sequencing (RADSeq) was then performed at the University of Wisconsin Biotechnology Center (UWBC, Madison, WI) using two enzymes (Pstl and Mspl) but barcoding only a single cut site (TGCA). 150 base-pair paired-end sequencing was then performed on an Illumina NovaSeq.

Raw Illumina reads were processed using iPyrad version 0.9 (*77*). We first demultiplexed the raw reads and trimmed low-quality base-pairs, and then aligned reads for each species to a reference genome. For *Hypocnemis*, *Phlegopsis*, and *Willisornis, we* used *Rhegmatorhina melanosticta* as a reference genome (*81*). For *Galbula*, *Malacoptila,* and *Thamnophilus*, we used *Galbula dea, Bucco capensis,* and *Sakesphorus luctuosus* as reference genomes, respectively (*82*). iPyrad uses bwa (*83*) to map cleaned illumina reads to a reference and determine homology among loci, and discards unmapped reads. We specifically applied a minimum coverage for statistical base-calling of six, with minimum trimmed read length of 35 bp, a maximum of 5% uncalled bases and 5% heterozygous sites, and a mapping threshold of 0.9. These steps were repeated to generate variant call format files for each species group, which were used and filtered for downstream analyses.

*Characterization of genomic variation*

We took multiple approaches to characterize genomic diversity across each species. First, we used iPyrad API analysis tools (*77*) to perform Principal Components Analysis (PCA) on each species. Because RADSeq datasets often contain a large proportion of missing data, we performed PCA after imputing missing haplotypes using k-means clustering implemented in the iPyrad API. This method allows imputation without *a priori* bias about population assignments. Specifically, we assumed k=8 for all species (except *Hypocnemis* because Inambari was not sampled) and used iterative clustering to group individuals into populations. For iterative clustering, we first sampled SNPs present across 90% of individuals in the assembly, then clustered with the assumed K. This was repeated five times, allowing more missing data at each successive iteration until reaching a minimum of 75% coverage at each sampled SNP. To impute, we then randomly sampled haplotypes based on the frequency of alleles within each of the populations defined by k-means. Because each species was characterized by a large amount of total genomic divergence across PC space, we split each species’ dataset into two population subsets based on phylogenetic relatedness (i.e., clades; see ASTRAL analysis below), and ran PCA on each data subset to improve visualization. We also utilized t-SNE, which is explained in further detail below. To evaluate the effects of missing data on the results, we further performed a PCA with k-means clustering on varying amounts of missing data from 55% to 95% coverage at each sampled SNP. This exercise showed that the results were robust to missing data thresholds and is available in our github repository at <https://github.com/lukemusher/Southern_Amazon_cophylogeography/tree/main/PCA_tSNE>.

As an independent method for assigning individuals to populations without *a priori* geographic bias, we used STRUCTURE v2.3.4 (*40*). We ran STRUCTURE on the entire dataset in addition to each of the two population subsets separately (see above), allowing for 50% missing individuals at each sampled SNP. We then performed 10 runs of 2.5 x 10^5^ generations with a burn-in of 5 x 10^4^ generations for each K-value of 2–12. Using clumpp (*84*), we optimally aligned these 10 runs for each K value and then chose the best value for K (the number of structured populations) for each species using the Evanno method, which chooses K based on the highest ΔK value (*85*).

We finally quantified genome-wide haplotype diversity (π), Watterson’s theta (θw), and Tajima’s D (D) in each of the eight *a priori* defined populations across all taxa using PopGenome (*86*). Moreover, we generated sliding windows of 5,000 bp with a jump of 2,500 bp and calculated π across the genomes of each population within each taxon. In a few instances, small sample size precluded estimation of some of these statistics.

*Environmental Niche Modeling*

To generate the ENMs for calculating least cost path distances, we downloaded locality data from the Global Biodiversity Information Facility for all six species. Because these species each have relatively large distributions that span most of Amazonia, and populations within each of these taxa is likely differentiated into multiple subspecies (if not full species), we then filtered these data to remove erroneous localities in addition to localities outside of our region of interest (southern Amazonia south of the Amazon river). We believe this should reduce noise and overfitting of the ENMs. We then thinned these data using the R package *spThin* *(87)* to reduce biases associated with sampling methods. We applied 10,000 replicates and retained five thinned datasets that maximized the number of filtered localities. We then randomly selected one thinned dataset to use for downstream analysis.

We created optimal ENMs using only climatic variables. We specifically used the 19 bioclimatic variables available on woldclim.org using a 30 arcsecond or ~1km resolution (*79*). For each species we sampled 10,000 points from a background defined as 10km larger than the minimum convex polygon of all GBIF occurrences. Background points were used for model tuning and final model creation.

We finally optimized ENMs for each species via a model tuning exercise, which varies regularization multipliers (RMs) and feature classes (FCs) for Maxent 3.4.1 using ENMeval (*88*, *89*). Varying RMs apply a weighting scheme for parameters included in the models while varying FCs allow for a range of response curves for model fitting. For each species we used a range of RMs from one to six in increments of 0.5 while also testing for the FCs “linear, “linear + quadratic”, “hinge”, “liner+quatratic+hinge”, and “linear+quatratic+hinge+product” at each RM increment. We used the “block” method of geographical partitioning to train and test datasets, which ensures generality and generates evaluation statistics for each model (*89*). To select the best model, we used sequential criteria to minimize overfitting and maximize the model’s discriminatory ability. First, we kept all models that displayed the lowest omission rate at the 10% omission threshold, which reduces overfitting (*90*). Second, we selected the model with the maximum test-AUC, which maximizes the model’s discriminatory ability to separate true presence localities from that of the background. We then projected these models to current climatic conditions.

*Phylogenomic trees and networks*

To characterize the evolutionary relationships among populations within each species, including potential reticulation, we employed three approaches. First, we estimated the historical relationships among populations using ASTRAL 5.7.3, (*41*). ASTRAL estimates the species tree from a set of unrooted gene trees under the multi-species coalescent model, with an assumption of no gene-flow among species. To estimate the gene trees for ASTRAL, we used RAxML 8.2.12 (*91*) on all loci greater than 400 bp in length. To reduce potential noise from gene tree estimation error, we then applied a filtering algorithm that identifies outlier loci (loci deviating from clock likeness expectations across the genome) (*61*). To assess robustness of the inferred population relationships, we also applied 100 bootstraps, subsampling gene trees only, using the *--gene-only* flag. We assigned all individuals in the ASTRAL analysis to each of the eight *a priori* defined populations allowing for alleles from multiple individuals per population.

To examine the evolutionary relationships within each species while relaxing the assumption of no gene-flow, we additionally applied Bayesian phylogenomic network analysis using the MCMC_GT algorithm implemented in PhyloNet (*44*). This algorithm uses a reverse-jump Markov Chain Monte Carlo (rjMCMC) to sample the posterior distribution of phylogenetic networks under a Multi-Species Network Coalescent model (MSNC). The MSNC models genome evolution as a network (as opposed to bifurcating tree), accounting for both incomplete lineage sorting and inter-lineage gene-flow (reticulation). We used the same set of gene trees estimated in RAxML and filtered under above criteria to estimate the network for each species. Gene trees were rooted based on the reference genome. We ran the rjMCMC for 5 x 10^7^ generations with a burnin of 5 x 10^6^, sampling every 10^5^ generations, using the pseudo-likelihood calculation, and allowing a maximum of two reticulate nodes per network (the rjMCMC will thus test networks with zero, one, and two reticulations only). The network topology with the highest posterior probability (PP) is chosen as the most credible network topology.

Finally, we used TreeMix 1.13 (*45*) to estimate the maximum likelihood population graph in the presence of gene-flow by attaching migration edges to a tree of bifurcating populations. Migration edges are assigned migration weights, which are related to the proportion of alleles in a population that are derived from migration. To examine the sensitivity of TreeMix output to missing data, we first ran TreeMix on subsamples of 50%, 60%, 70%, 80%, and 90% missing data with one migration edge (m=1). We then examined the model fit for varying values of m (m=0– 3) with zero representing pure isolation. To do so, we performed 250 TreeMix replicates, randomly choosing missing data thresholds between 50% and 95% and quantifying the likelihood of the model and averaging across all replicates. This was repeated for each value of m.

Because unstructured populations may experience high gene-flow, for both PhyloNet and TreeMix we assigned individuals to populations based on their assignments in STRUCTURE. However, because STRUCTURE analyses resulted in underestimation of population structure for *Thamnophilus* and *Phlegopsis* when compared with PCA and previous work (*8*, *24*), we split one of the population assignments in two based on PCA results with the goal of better understanding spatial patterns of isolation and reticulation in these groups.

*Demographic Modeling*

To estimate divergence times and effective population sizes in the presence of gene-flow, we applied a Bayesian approach using Generalized Phylogenetic Coalescent Sampler (G-PhoCS) (*47*). G-PhoCS integrates full likelihood computation across all possible gametic phasings based on unphased loci and a predefined demographic model to estimate mutation-scaled effective population sizes (θ) and divergence times (τ) for all populations. The software also allows estimation of continuous gene-flow between sets of current or ancestral populations. We constructed demographic models for each species-group based on the results from PhyloNet. Specifically, reticulate edges in PhyloNet with higher inheritance probabilities (>0.5) were treated as population divergences, whereas the edges with lower probabilities (<0.5), were treated as migration edges. We allowed for all possible migration edges based on both PhyloNet and TreeMix results. We applied priors on root τ (α = 3.0; β= 1,000) and θ (α = 5.0; β= 1000), and for each population and divergence (τ α = 1.0, β= 50,000; θ α = 5.0, β= 1,000). We then ran the MCMC for 5 x 10^5^ iterations after a burn-in of ten percent, sampling every ten iterations.

To convert τ and θ outputs to absolute divergence times (T) and effective population sizes (Ne), respectively, we used the formulas Ne = θ/4µ and T = τG/µ, where µ is the mutation rate in substitutions per site per generation and G is the generation time in years. Because mutation rates are known to vary significantly across avian lineages, we estimated T and Ne while allowing for some variance in µ (*92*, *93*). To do so, we generated a normal distribution of values for µ with mean = 4.6 x 10^-9^and standard deviation = 10^-9^. Although standard deviation in µ across the tree of birds may be much higher than this, it offers a reasonable assessment of uncertainty around our inferred divergence dates. We then performed 100,000 replicates, randomly sampling values of µ from the normal distribution and values of τ from the posterior distribution of the G-PhoCS analysis. We repeated this for values of theta to estimate Ne. This resulted in distributions of T and Ne for each divergence event and population size, respectively. Mean µ and G=2 were assumed based on previous work (*94*). We then plotted mean values for T with 95% confidence intervals in R. Though interpopulation migration was estimated, its main function in the model was to improve estimates of T and Ne for each group. To assess the robustness of our inferred divergence times, we also compared these results to divergence times inferred from Sanger sequencing data in another study (*8*).

**Results**

*Structure Analysis*

The best value of k ranged from three to five for all datasets (Figure S1). Results for the split datasets showed finer scale structure than the complete datasets, though still not as fine as PCA and t-SNE. However, the structure results recapitulated the general pattern of population structure associated with rivers. The results of the structure analyses can be found in Figure S1.

*Population genetic summary statistics*

To characterize population genomic variation in each species, we also estimated a series of genome-wide summary statistics (Table S2). Among species, genome-wide nucleotide diversity (π) ranged from very low (*Thamnophilus*) to quite high (*Malacoptila*), but among populations within each species π varied relatively little. Exceptions to the latter pattern include *Malacoptila*, where West Inambari and Rondonia 1 populations showed lower overall π, and *Phlegopsis* where East and West Inambari showed higher π. Values of Watterson’s theta (θ_w_) tended to be somewhat higher than values of π within each population, indicating a slight excess of rare alleles in most populations. These statistics were concordant with estimates of Tajima’s D (D), which were negative for all populations. Most values of D were close to zero, though, with the lowest values being -1.7 in the Rondonia 5 population of *T. aethiops*, suggesting recent expansions or bottlenecks are unlikely to have occurred in these populations (*95*). Due to the nature of independently assembled RADseq datasets, it may not be appropriate to directly compare results between species, so we would exercise caution in the interpretation of these results.

*Phylonet*

For four out of the six species, the most credible network was recovered with high posterior support (PP=1.0) in PhyloNet (*44*). However, for two groups, *Malacoptila* and *Thamnophilus*, the most credible topology was less supported. The topology for *Malacoptila* was recovered with PP=0.76, while *Thamnophilus* was recovered with PP=0.67 (Figure S6). These networks were nearly identical to the most credible network topologies shown in Figure 5 of the main text, but differ only in the location of reticulate parent nodes deep in each network (as opposed to at the tips).

**Supplemental Tables:**

| **Institution** | **Tissue_No** | **CollectNo** | **amcc** | **genus** | **species** | **lat** | **long** |
| --- | --- | --- | --- | --- | --- | --- | --- |
| MPEG | 82508 | PUR025 |  | *Galbula* | *cyanicollis* | -7.6945 | -65.2235 |
| MPEG | T12275 | TUP011 |  | *Galbula* | *cyanicollis* | -4.0833 | -60.6605 |
| MPEG | T12279 | TUP015 |  | *Galbula* | *cyanicollis* | -4.0833 | -60.6605 |
| MPEG | T12392 | TUP050 |  | *Galbula* | *cyanicollis* | -4.0833 | -60.6605 |
| MPEG | T13184 | OM133 |  | *Galbula* | *cyanicollis* | -7.5183 | -63.3378 |
| MPEG | T26228 | PUR024 |  | *Galbula* | *cyanicollis* | -7.6950 | -65.2241 |
| MPEG | T26229 | PUR025 |  | *Galbula* | *cyanicollis* | -7.6950 | -65.2241 |
| MPEG | T26252 | PUR048 |  | *Galbula* | *cyanicollis* | -7.6950 | -65.2241 |
| MPEG | T10897 | MPDS1229 |  | *Galbula* | *cyanicollis* | -6.5190 | -57.4446 |
| MPEG | T11062 | PIME362 |  | *Galbula* | *cyanicollis* | -5.0381 | -55.5189 |
| MPEG | T16771 | PNSP054 |  | *Galbula* | *cyanicollis* | -5.8778 | -52.7954 |
| MPEG | T1705 | UHE410 |  | *Galbula* | *cyanicollis* | -3.5298 | -51.7329 |
| MPEG | T18744 | JAT(B)012 |  | *Galbula* | *cyanicollis* | -5.5695 | -57.1281 |
| MPEG | T19429 | JAT(C)262 |  | *Galbula* | *cyanicollis* | -5.2303 | -56.9322 |
| MPEG | T19520 | JAT(A)261 |  | *Galbula* | *cyanicollis* | -6.1102 | -57.6130 |
| MPEG | T19765 | JAT(A)474 |  | *Galbula* | *cyanicollis* | -5.7929 | -57.4000 |
| MPEG | T2497 | BR163-159 |  | *Galbula* | *cyanicollis* | -7.1914 | -55.4954 |
| MPEG | T6579 | TM051 |  | *Galbula* | *cyanicollis* | -5.6500 | -55.5167 |
| MPEG | T9133 | FLJA055 |  | *Galbula* | *cyanicollis* | -7.1351 | -55.4817 |
| MPEG | T3343 | CUJ086 |  | *Galbula* | *cyanicollis* | -10.8333 | -64.7500 |
| MPEG | T3384 | CUJ142 |  | *Galbula* | *cyanicollis* | -10.8333 | -64.7500 |
| MPEG | T3385 | CUJ143 |  | *Galbula* | *cyanicollis* | -10.8333 | -64.7500 |
| MPEG | J296 | LJM220 |  | *Galbula* | *cyanicollis* | -7.5672 | -60.6882 |
| MPEG | J477 | LJM264 |  | *Galbula* | *cyanicollis* | -7.6951 | -60.9124 |
| AMNH | J773 | GDR403 | 261446 | *Galbula* | *cyanicollis* | -7.5672 | -60.6882 |
| MPEG | T13251 | OM201 |  | *Galbula* | *cyanicollis* | -8.9091 | -62.0001 |
| MPEG | T363 | MAR054 |  | *Galbula* | *cyanicollis* | -8.6873 | -61.4082 |
| MPEG | T364 | MAR055 |  | *Galbula* | *cyanicollis* | -8.6873 | -61.4082 |
| AMNH | J691 | LJM311 | 261356 | *Galbula* | *cyanicollis* | -8.1776 | -60.4531 |
| AMNH | J694 | GDR375 | 261360 | *Galbula* | *cyanicollis* | -8.1776 | -60.4531 |
| LSUMNS | 80582 |  |  | *Galbula* | *cyanicollis* | -5.2889 | -59.6936 |
| LSUMNS | 80701 |  |  | *Galbula* | *cyanicollis* | -5.2889 | -59.6936 |
| LSUMNS | 80801 |  |  | *Galbula* | *cyanicollis* | -5.2518 | -59.6947 |
| LSUMNS | 80826 |  |  | *Galbula* | *cyanicollis* | -5.2889 | -59.6936 |
| LSUMNS | 81108 |  |  | *Galbula* | *cyanicollis* | -5.7991 | -59.2686 |
| LSUMNS | 81118 |  |  | *Galbula* | *cyanicollis* | -5.7991 | -59.2686 |
| LSUMNS | 85499 |  |  | *Galbula* | *cyanicollis* | -5.9333 | -59.1761 |
| LSUMNS | 85678 |  |  | *Galbula* | *cyanicollis* | -5.9506 | -59.1739 |
| LSUMNS | 85356 |  |  | *Galbula* | *cyanicollis* | -5.9444 | -59.1639 |
| LSUMNS | 86297 |  |  | *Galbula* | *cyanicollis* | -5.2619 | -59.0681 |
| LSUMNS | 86321 |  |  | *Galbula* | *cyanicollis* | -6.2519 | -59.0681 |
| LSUMNS | 86458 |  |  | *Galbula* | *cyanicollis* | -6.2519 | -59.0681 |
| LSUMNS | 86478 |  |  | *Galbula* | *cyanicollis* | -6.2619 | -59.0681 |
| MPEG | T14558 | ARA032 |  | *Galbula* | *cyanicollis* | -3.1500 | -55.8167 |
| MPEG | T16693 | ARAII091 |  | *Galbula* | *cyanicollis* | -2.5868 | -55.1952 |
| MPEG | T18563 | JAT(C)144 |  | *Galbula* | *cyanicollis* | -5.0733 | -56.8559 |
| MPEG | T18620 | JAT(A)023 |  | *Galbula* | *cyanicollis* | -5.7385 | -57.3555 |
| MPEG | T12385 | JUR040 |  | *Galbula* | *cyanicollis* | -7.9600 | -69.9458 |
| MPEG | T23196 | JUT011 |  | *Galbula* | *cyanicollis* | -3.2711 | -67.3252 |
| MPEG | T310 | PUC270 |  | *Galbula* | *cyanicollis* | -4.8732 | -65.3026 |
| MPEG | T7636 | CUJ191 |  | *Galbula* | *cyanicollis* | -4.9347 | -68.1725 |
| INPA | A311 |  |  | *Hypocnemis* | *cantator* | -8.8305 | -63.9605 |
| MPEG | T355 | MAR046 |  | *Hypocnemis* | *cantator* | -10.8333 | -64.7500 |
| INPA | A409 |  |  | *Hypocnemis* | *cantator* | -6.3000 | -60.3333 |
| INPA | A410 |  |  | *Hypocnemis* | *cantator* | -6.3000 | -60.3333 |
| INPA | A521 |  |  | *Hypocnemis* | *cantator* | -6.3000 | -60.3333 |
| INPA | A547 |  |  | *Hypocnemis* | *cantator* | -6.2500 | -60.4500 |
| INPA | A551 |  |  | *Hypocnemis* | *cantator* | -6.3000 | -60.3333 |
| MPEG | T1842 | MPDS659 |  | *Hypocnemis* | *cantator* | -7.4667 | -62.8167 |
| MPEG | T366 | MAR059 |  | *Hypocnemis* | *cantator* | -8.6873 | -61.4082 |
| MPEG | T385 | MAR081 |  | *Hypocnemis* | *cantator* | -8.6957 | -61.4082 |
| MPEG | T471 | MAR182 |  | *Hypocnemis* | *cantator* | -8.6504 | -61.4201 |
| INPA | A272 |  |  | *Hypocnemis* | *cantator* | -6.0167 | -60.1667 |
| INPA | A273 |  |  | *Hypocnemis* | *cantator* | -6.0167 | -60.1667 |
| MPEG | T7114 | AMANA148 |  | *Hypocnemis* | *cantator* | -5.9035 | -57.6917 |
| MPEG | T15847 | MAD023 |  | *Hypocnemis* | *ochrogyna* | -8.2239 | -63.1818 |
| INPA | A24346 |  |  | *Hypocnemis* | *peruviana* | -7.8319 | -63.8448 |
| MPEG | J213 | MAR091 |  | *Hypocnemis* | *rondoni* | -7.5672 | -60.6882 |
| MPEG | J305 | GDR249 |  | *Hypocnemis* | *rondoni* | -7.5672 | -60.6882 |
| MPEG | J508 | GT108 |  | *Hypocnemis* | *rondoni* | -7.6951 | -60.9124 |
| AMNH | J774 | LJM325 | 261438 | *Hypocnemis* | *rondoni* | -7.5672 | -60.6882 |
| AMNH | J775 | LJM326 | 261449 | *Hypocnemis* | *rondoni* | -7.5672 | -60.6882 |
| AMNH | J796 | GDR411 | 261467 | *Hypocnemis* | *rondoni* | -7.5672 | -60.6882 |
| INPA | A11597 |  |  | *Hypocnemis* | *striata* | -4.7451 | -56.4127 |
| INPA | A15208 |  |  | *Hypocnemis* | *striata* | -3.3500 | -55.2000 |
| INPA | A15210 |  |  | *Hypocnemis* | *striata* | -3.3500 | -55.2000 |
| INPA | A16571 |  |  | *Hypocnemis* | *striata* | -5.0833 | -56.4333 |
| INPA | A7597 |  |  | *Hypocnemis* | *striata* | -4.7811 | -56.4598 |
| INPA | A9955 |  |  | *Hypocnemis* | *striata* | -4.7667 | -56.5833 |
| MPEG | T16744 | PNSP028 |  | *Hypocnemis* | *striata* | -5.8778 | -52.7954 |
| MPEG | T17858 | PPS146 |  | *Hypocnemis* | *striata* | -9.9381 | -54.3416 |
| MPEG | J364 | GT067 |  | *Hypocnemis* | *striata* | -7.6998 | -60.8905 |
| MPEG | J368 | GT068 |  | *Hypocnemis* | *striata* | -7.6998 | -60.8905 |
| MPEG | J370 | GT069 |  | *Hypocnemis* | *striata* | -7.6998 | -60.8905 |
| MPEG | J374 | GDR267 |  | *Hypocnemis* | *striata* | -7.6998 | -60.8905 |
| MPEG | J408 | LJM244 |  | *Hypocnemis* | *striata* | -7.6998 | -60.8905 |
| AMNH | J621 | GDR351 | 261288 | *Hypocnemis* | *striata* | -8.1776 | -60.4531 |
| AMNH | J664 | LJM304 | 261328 | *Hypocnemis* | *striata* | -8.1776 | -60.4531 |
| AMNH | J665 | LJM305 | 261329 | *Hypocnemis* | *striata* | -8.1776 | -60.4531 |
| AMNH | J711 | GDR380 | 261376 | *Hypocnemis* | *striata* | -8.1776 | -60.4531 |
| AMNH | J762 | MAR149 | 261432 | *Hypocnemis* | *striata* | -8.1776 | -60.4531 |
| AMNH | J765 | LJM322 | 261433 | *Hypocnemis* | *striata* | -8.1776 | -60.4531 |
| LSUMNS | 77860 |  |  | *Hypocnemis* | *striata* | -7.3475 | -58.1572 |
| LSUMNS | 78249 |  |  | *Hypocnemis* | *striata* | -7.5133 | -58.2536 |
| LSUMNS | 80727 |  |  | *Hypocnemis* | *striata* | -5.2889 | -59.6936 |
| LSUMNS | 80800 |  |  | *Hypocnemis* | *striata* | -5.2518 | -59.6947 |
| LSUMNS | 81143 |  |  | *Hypocnemis* | *striata* | -5.8025 | -59.2563 |
| LSUMNS | 85680 |  |  | *Hypocnemis* | *striata* | -5.9506 | -59.1739 |
| LSUMNS | 85970 |  |  | *Hypocnemis* | *striata* | -6.7056 | -59.0789 |
| AMNH | J525 | GDR317 | 261190 | *Hypocnemis* | *striata* | -7.7037 | -60.5957 |
| AMNH | J530 | GDR319 | 261196 | *Hypocnemis* | *striata* | -7.7037 | -60.5957 |
| AMNH | J536 | GT116 | 261202 | *Hypocnemis* | *striata* | -7.7037 | -60.5957 |
| AMNH | J572 | LJM285 | 261237 | *Hypocnemis* | *striata* | -7.7037 | -60.5957 |
| LSUMNS | 81279 |  |  | *Hypocnemis* | *striata* | -4.0508 | -59.1153 |
| LSUMNS | 86405 |  |  | *Hypocnemis* | *striata* | -5.2619 | -59.0681 |
| INPA | A14546 |  |  | *Hypocnemis* | *striata* | -5.0622 | -56.8739 |
| INPA | A4899 |  |  | *Hypocnemis* | *striata* | -4.6731 | -56.6231 |
| MPEG | T10207 | FPR071 |  | *Hypocnemis* | *striata* | -3.9070 | -58.4018 |
| MPEG | T11900 | PIME197 |  | *Hypocnemis* | *striata* | -5.5711 | -57.3064 |
| MPEG | T12194 | MAM007 |  | *Hypocnemis* | *striata* | -3.3279 | -56.3635 |
| MPEG | T24564 | ALC017 |  | *Hypocnemis* | *striata* | -2.7833 | -56.0000 |
| INPA | A10311 |  |  | *Malacoptila* | *rufa* | -4.9833 | -61.5667 |
| INPA | A10329 |  |  | *Malacoptila* | *rufa* | -4.9833 | -61.5667 |
| INPA | A1380 |  |  | *Malacoptila* | *rufa* | -4.1500 | -60.1333 |
| INPA | A2741 |  |  | *Malacoptila* | *rufa* | -4.4167 | -60.9167 |
| INPA | A436 |  |  | *Malacoptila* | *rufa* | -5.2417 | -60.7167 |
| INPA | A440 |  |  | *Malacoptila* | *rufa* | -5.2417 | -60.7167 |
| INPA | A16195 |  |  | *Malacoptila* | *rufa* | -4.8815 | -56.4462 |
| INPA | A9235 |  |  | *Malacoptila* | *rufa* | -5.0833 | -56.4333 |
| INPA | A9903 |  |  | *Malacoptila* | *rufa* | -4.7167 | -56.4333 |
| MPEG | T11079 | PIME380 |  | *Malacoptila* | *rufa* | -5.0381 | -55.5189 |
| MPEG | T11238 | PIME276 |  | *Malacoptila* | *rufa* | -4.5408 | -55.2106 |
| MPEG | T12541 | PIME385 |  | *Malacoptila* | *rufa* | -7.1292 | -55.4252 |
| MPEG | T1649 | WM385 |  | *Malacoptila* | *rufa* | -3.3561 | -54.9492 |
| MPEG | T16553 | MF014 |  | *Malacoptila* | *rufa* | -4.4689 | -55.8503 |
| MPEG | T19782 | JAT(C)325 |  | *Malacoptila* | *rufa* | -5.4307 | -56.9046 |
| MPEG | T6500 | TM013 |  | *Malacoptila* | *rufa* | -6.0333 | -55.3167 |
| MPEG | T6577 | TM049 |  | *Malacoptila* | *rufa* | -5.6500 | -55.5167 |
| INPA | A207 |  |  | *Malacoptila* | *rufa* | -9.5285 | -65.3007 |
| MPEG | T3228 | OP098 |  | *Malacoptila* | *rufa* | -10.8333 | -64.7500 |
| MPEG | T7634 | OP097 |  | *Malacoptila* | *rufa* | -10.8333 | -64.7500 |
| INPA | A474 |  |  | *Malacoptila* | *rufa* | -6.3000 | -60.4000 |
| MPEG | J265 | LJM215 |  | *Malacoptila* | *rufa* | -7.5672 | -60.6882 |
| MPEG | T13253 | OM203 |  | *Malacoptila* | *rufa* | -8.9091 | -62.0001 |
| MPEG | T3164 | MPDS759 |  | *Malacoptila* | *rufa* | -6.5500 | -62.0500 |
| MPEG | T368 | MAR063 |  | *Malacoptila* | *rufa* | -8.6873 | -61.4082 |
| MPEG | T381 | MAR076 |  | *Malacoptila* | *rufa* | -8.6957 | -61.4082 |
| MPEG | T476 | MAR187 |  | *Malacoptila* | *rufa* | -8.6504 | -61.4201 |
| MPEG | T494 | MAR218 |  | *Malacoptila* | *rufa* | -8.6504 | -61.4201 |
| AMNH | J640 | LJM300 | 261306 | *Malacoptila* | *rufa* | -8.1776 | -60.4531 |
| AMNH | J676 | LJM307 | 261342 | *Malacoptila* | *rufa* | -8.1776 | -60.4531 |
| LSUMNS | 80819 |  |  | *Malacoptila* | *rufa* | -5.8025 | -59.2563 |
| LSUMNS | 81347 |  |  | *Malacoptila* | *rufa* | -4.0508 | -59.1153 |
| LSUMNS | 85426 |  |  | *Malacoptila* | *rufa* | -5.9506 | -59.1739 |
| LSUMNS | 77750 |  |  | *Malacoptila* | *rufa* | -7.3475 | -58.1572 |
| LSUMNS | 78182 |  |  | *Malacoptila* | *rufa* | -7.5133 | -58.2536 |
| LSUMNS | 85919 |  |  | *Malacoptila* | *rufa* | -6.7556 | -59.0756 |
| INPA | A11834 |  |  | *Malacoptila* | *rufa* | -4.6935 | -56.6385 |
| INPA | A15176 |  |  | *Malacoptila* | *rufa* | -3.3167 | -55.3333 |
| INPA | A5487 |  |  | *Malacoptila* | *rufa* | -4.4950 | -56.3003 |
| MPEG | T10184 | FPR048 |  | *Malacoptila* | *rufa* | -3.9461 | -58.4561 |
| MPEG | T11780 | PIME152 |  | *Malacoptila* | *rufa* | -6.2469 | -57.8750 |
| MPEG | T14532 | ARA006 |  | *Malacoptila* | *rufa* | -3.1500 | -55.8167 |
| MPEG | T14622 | ARA096 |  | *Malacoptila* | *rufa* | -2.7833 | -55.6000 |
| MPEG | T753 | JRT100 |  | *Malacoptila* | *rufa* | -2.4667 | -56.0000 |
| INPA | A7875 |  |  | *Malacoptila* | *rufa* | -4.9833 | -62.9667 |
| MPEG | T14456 | AMA610 |  | *Malacoptila* | *rufa* | -4.5303 | -71.6163 |
| MPEG | T23245 | JUT060 |  | *Malacoptila* | *rufa* | -3.2711 | -67.3252 |
| MPEG | T23246 | JUT061 |  | *Malacoptila* | *rufa* | -3.2711 | -67.3252 |
| MPEG | T23416 | JUT238 |  | *Malacoptila* | *rufa* | -3.2235 | -67.4418 |
| MPEG | T23478 | JUT303 |  | *Malacoptila* | *rufa* | -3.2233 | -67.4417 |
| LSUMNS | 80034 |  |  | *Phlegopsis* | *nigromaculata* | -10.5400 | -66.0700 |
| MPEG | T15938 | MAD116 |  | *Phlegopsis* | *nigromaculata* | -7.7760 | -62.9414 |
| MPEG | T3609 | UFAC217 |  | *Phlegopsis* | *nigromaculata* | -10.1284 | -67.3285 |
| MPEG | T3611 | UFAC219 |  | *Phlegopsis* | *nigromaculata* | -10.1284 | -67.3285 |
| MPEG | T3817 | UFAC592 |  | *Phlegopsis* | *nigromaculata* | -9.3262 | -68.3356 |
| MPEG | T4043 | UFAC081 |  | *Phlegopsis* | *nigromaculata* | -9.7567 | -67.6706 |
| MPEG | T4051 | UFAC090 |  | *Phlegopsis* | *nigromaculata* | -9.7567 | -67.6706 |
| MPEG | T4313 | UFAC907 |  | *Phlegopsis* | *nigromaculata* | -9.9006 | -68.4756 |
| MPEG | T4404 | UFAC858 |  | *Phlegopsis* | *nigromaculata* | -9.9006 | -68.4756 |
| MPEG | T5850 | UFAC1725 |  | *Phlegopsis* | *nigromaculata* | -10.6368 | -67.8155 |
| MPEG | T5940 | UFAC1202 |  | *Phlegopsis* | *nigromaculata* | -9.9590 | -67.7326 |
| MPEG | T5974 | UFAC1445 |  | *Phlegopsis* | *nigromaculata* | -9.1573 | -69.7634 |
| INPA | A14342 |  |  | *Phlegopsis* | *nigromaculata* | -4.8875 | -56.4242 |
| INPA | A15277 |  |  | *Phlegopsis* | *nigromaculata* | -3.3500 | -55.2000 |
| INPA | A7066 |  |  | *Phlegopsis* | *nigromaculata* | -5.0667 | -56.4333 |
| MPEG | T10673 | PIME117 |  | *Phlegopsis* | *nigromaculata* | -3.4729 | -54.5655 |
| MPEG | T10940 | MPDS1309 |  | *Phlegopsis* | *nigromaculata* | -6.5190 | -57.4446 |
| MPEG | T11193 | PIME229 |  | *Phlegopsis* | *nigromaculata* | -4.6088 | -55.4966 |
| MPEG | T11222 | PIME259 |  | *Phlegopsis* | *nigromaculata* | -4.5408 | -55.2106 |
| MPEG | T12345 | FAL021 |  | *Phlegopsis* | *nigromaculata* | -5.4913 | -55.1383 |
| MPEG | T12854 | PIME531 |  | *Phlegopsis* | *nigromaculata* | -7.1296 | -55.7174 |
| MPEG | T1642 | WM366 |  | *Phlegopsis* | *nigromaculata* | -3.3561 | -54.9492 |
| MPEG | T18703 | JAT(A)107 |  | *Phlegopsis* | *nigromaculata* | -5.7668 | -57.2876 |
| INPA | A3255 |  |  | *Phlegopsis* | *nigromaculata* | -9.2668 | -64.4001 |
| MPEG | T15863 | MAD039 |  | *Phlegopsis* | *nigromaculata* | -8.2239 | -63.1818 |
| MPEG | T15868 | MAD044 |  | *Phlegopsis* | *nigromaculata* | -8.2239 | -63.1818 |
| MPEG | T15871 | MAD047 |  | *Phlegopsis* | *nigromaculata* | -8.2239 | -63.1818 |
| MPEG | T22153 | MSF415 |  | *Phlegopsis* | *nigromaculata* | -10.9112 | -65.2240 |
| MPEG | T3261 | OP151 |  | *Phlegopsis* | *nigromaculata* | -10.8333 | -64.7500 |
| INPA | A2418 |  |  | *Phlegopsis* | *nigromaculata* | -9.4422 | -61.6838 |
| INPA | A542 |  |  | *Phlegopsis* | *nigromaculata* | -6.2500 | -60.4500 |
| MPEG | J210 | GT017 |  | *Phlegopsis* | *nigromaculata* | -7.5672 | -60.6882 |
| MPEG | J227 | GDR226 |  | *Phlegopsis* | *nigromaculata* | -7.5672 | -60.6882 |
| MPEG | J260 | LJM214 |  | *Phlegopsis* | *nigromaculata* | -7.5672 | -60.6882 |
| MPEG | J434 | GDR287 |  | *Phlegopsis* | *nigromaculata* | -7.6951 | -60.9124 |
| MPEG | J461 | MAR115 |  | *Phlegopsis* | *nigromaculata* | -7.6951 | -60.9124 |
| MPEG | J462 | GT095 |  | *Phlegopsis* | *nigromaculata* | -7.6951 | -60.9124 |
| MPEG | J485 | GT102 |  | *Phlegopsis* | *nigromaculata* | -7.6951 | -60.9124 |
| MPEG | T369 | MAR064 |  | *Phlegopsis* | *nigromaculata* | -8.6873 | -61.4082 |
| MPEG | T443 | MAR147 |  | *Phlegopsis* | *nigromaculata* | -8.6957 | -61.4082 |
| MPEG | T467 | MAR176 |  | *Phlegopsis* | *nigromaculata* | -8.6504 | -61.4201 |
| MPEG | J361 | GDR263 |  | *Phlegopsis* | *nigromaculata* | -7.6998 | -60.8905 |
| MPEG | J363 | LJM234 |  | *Phlegopsis* | *nigromaculata* | -7.6998 | -60.8905 |
| MPEG | J371 | GDR266 |  | *Phlegopsis* | *nigromaculata* | -7.6998 | -60.8905 |
| MPEG | J373 | GT070 |  | *Phlegopsis* | *nigromaculata* | -7.6998 | -60.8905 |
| MPEG | J381 | GDR269 |  | *Phlegopsis* | *nigromaculata* | -7.6998 | -60.8905 |
| MPEG | J385 | GT074 |  | *Phlegopsis* | *nigromaculata* | -7.6998 | -60.8905 |
| MPEG | J389 | MAR107 |  | *Phlegopsis* | *nigromaculata* | -7.6998 | -60.8905 |
| MPEG | J417 | MAR110 |  | *Phlegopsis* | *nigromaculata* | -7.6998 | -60.8905 |
| AMNH | J684 | LJM310 | 261350 | *Phlegopsis* | *nigromaculata* | -8.1776 | -60.4531 |
| AMNH | J724 | LJM317 | 261392 | *Phlegopsis* | *nigromaculata* | -8.1776 | -60.4531 |
| LSUMNS | 80555 |  |  | *Phlegopsis* | *nigromaculata* | -5.2803 | -59.6959 |
| LSUMNS | 80684 |  |  | *Phlegopsis* | *nigromaculata* | -5.2889 | -59.6936 |
| LSUMNS | 80802 |  |  | *Phlegopsis* | *nigromaculata* | -5.2518 | -59.6947 |
| LSUMNS | 80874 |  |  | *Phlegopsis* | *nigromaculata* | -5.8025 | -59.2563 |
| LSUMNS | 85430 |  |  | *Phlegopsis* | *nigromaculata* | -5.9506 | -59.1739 |
| LSUMNS | 86072 |  |  | *Phlegopsis* | *nigromaculata* | -6.7547 | -59.0831 |
| AMNH | J551 | LJM282 | 261217 | *Phlegopsis* | *nigromaculata* | -7.7037 | -60.5957 |
| AMNH | J602 | GDR344 | 261267 | *Phlegopsis* | *nigromaculata* | -7.7037 | -60.5957 |
| AMNH | J603 | GT140 | 261269 | *Phlegopsis* | *nigromaculata* | -7.7037 | -60.5957 |
| AMNH | J614 | GDR350 | 261280 | *Phlegopsis* | *nigromaculata* | -7.7037 | -60.5957 |
| AMNH | J617 | GT142 | 261281 | *Phlegopsis* | *nigromaculata* | -7.7037 | -60.5957 |
| LSUMNS | 77876 |  |  | *Phlegopsis* | *nigromaculata* | -7.3475 | -58.1572 |
| LSUMNS | 78155 |  |  | *Phlegopsis* | *nigromaculata* | -7.5133 | -58.2536 |
| LSUMNS | 85721 |  |  | *Phlegopsis* | *nigromaculata* | -5.7979 | -59.2308 |
| INPA | A15120 |  |  | *Phlegopsis* | *nigromaculata* | -3.3167 | -55.3333 |
| MPEG | T10204 | FPR068 |  | *Phlegopsis* | *nigromaculata* | -3.9070 | -58.4018 |
| MPEG | T10967 | MPDS0812 |  | *Phlegopsis* | *nigromaculata* | -2.6000 | -56.1833 |
| MPEG | T11888 | PIME181 |  | *Phlegopsis* | *nigromaculata* | -5.8976 | -57.6966 |
| MPEG | T14543 | ARA017 |  | *Phlegopsis* | *nigromaculata* | -3.1500 | -55.8167 |
| MPEG | T16698 | ARAII096 |  | *Phlegopsis* | *nigromaculata* | -2.5868 | -55.1952 |
| MPEG | T9076 | PIME030 |  | *Phlegopsis* | *nigromaculata* | -2.3983 | -55.7884 |
| INPA | A7862 |  |  | *Phlegopsis* | *nigromaculata* | -4.9833 | -62.9667 |
| INPA | A7911 |  |  | *Phlegopsis* | *nigromaculata* | -4.9833 | -62.9667 |
| INPA | A7928 |  |  | *Phlegopsis* | *nigromaculata* | -4.9833 | -62.9667 |
| MPEG | T6243 | UFAC1864 |  | *Phlegopsis* | *nigromaculata* | -8.4599 | -70.5564 |
| MPEG | 82491 | PUR008 |  | *Thamnophilus* | *aethiops* | -7.6945 | -65.2235 |
| MPEG | 82606 | PUR124 |  | *Thamnophilus* | *aethiops* | -7.3365 | -65.1600 |
| MPEG | 82608 | PUR126 |  | *Thamnophilus* | *aethiops* | -7.3365 | -65.1600 |
| INPA | A21562 |  |  | *Thamnophilus* | *aethiops* | -5.2654 | -61.9444 |
| INPA | A2720 |  |  | *Thamnophilus* | *aethiops* | -3.6833 | -60.3167 |
| INPA | A2833 |  |  | *Thamnophilus* | *aethiops* | -4.9833 | -61.5667 |
| INPA | A3559 |  |  | *Thamnophilus* | *aethiops* | -8.8409 | -64.0616 |
| INPA | A439 |  |  | *Thamnophilus* | *aethiops* | -5.2417 | -60.7167 |
| MPEG | T12313 | TUP057 |  | *Thamnophilus* | *aethiops* | -4.0833 | -60.6605 |
| MPEG | T13219 | OM168 |  | *Thamnophilus* | *aethiops* | -7.5183 | -63.3378 |
| INPA | A15279 |  |  | *Thamnophilus* | *aethiops* | -3.3500 | -55.2000 |
| MPEG | T10679 | PIME123 |  | *Thamnophilus* | *aethiops* | -3.4729 | -54.5655 |
| MPEG | T13575 | MIRI020 |  | *Thamnophilus* | *aethiops* | -4.2833 | -55.9167 |
| MPEG | T17307 | PRTS003 |  | *Thamnophilus* | *aethiops* | -5.9560 | -56.4488 |
| MPEG | T19420 | JAT(C)252 |  | *Thamnophilus* | *aethiops* | -5.2303 | -56.9322 |
| MPEG | T8268 | TM006 |  | *Thamnophilus* | *aethiops* | -6.0667 | -55.3167 |
| INPA | A317 |  |  | *Thamnophilus* | *aethiops* | -8.8533 | -63.8630 |
| INPA | A324 |  |  | *Thamnophilus* | *aethiops* | -8.8533 | -63.8630 |
| MPEG | T3237 | OP115 |  | *Thamnophilus* | *aethiops* | -10.8333 | -64.7500 |
| MPEG | T3260 | OP150 |  | *Thamnophilus* | *aethiops* | -10.8333 | -64.7500 |
| MPEG | T3376 | CUJ133 |  | *Thamnophilus* | *aethiops* | -10.8333 | -64.7500 |
| INPA | A458 |  |  | *Thamnophilus* | *aethiops* | -6.3000 | -60.4000 |
| INPA | A509 |  |  | *Thamnophilus* | *aethiops* | -6.3000 | -60.4000 |
| INPA | A522 |  |  | *Thamnophilus* | *aethiops* | -6.3000 | -60.3333 |
| MPEG | J249 | GDR233 |  | *Thamnophilus* | *aethiops* | -7.5672 | -60.6882 |
| MPEG | J252 | GDR234 |  | *Thamnophilus* | *aethiops* | -7.5672 | -60.6882 |
| MPEG | J261 | ROS006 |  | *Thamnophilus* | *aethiops* | -7.5672 | -60.6882 |
| MPEG | J298 | GDR247 |  | *Thamnophilus* | *aethiops* | -7.5672 | -60.6882 |
| MPEG | T13278 | OM228 |  | *Thamnophilus* | *aethiops* | -8.9091 | -62.0000 |
| MPEG | T2166 | MPDS665 |  | *Thamnophilus* | *aethiops* | -7.4667 | -62.8167 |
| MPEG | T2207 | MPDS720 |  | *Thamnophilus* | *aethiops* | -7.5500 | -62.5500 |
| MPEG | T4355 | MPDS721 |  | *Thamnophilus* | *aethiops* | -7.5500 | -62.5500 |
| MPEG | J319 | LJM225 |  | *Thamnophilus* | *aethiops* | -7.6998 | -60.8905 |
| MPEG | J419 | GDR283 |  | *Thamnophilus* | *aethiops* | -7.6998 | -60.8905 |
| AMNH | J678 | GDR370 | 261344 | *Thamnophilus* | *aethiops* | -8.1776 | -60.4531 |
| AMNH | J683 | GT167 | 261349 | *Thamnophilus* | *aethiops* | -8.1776 | -60.4531 |
| LSUMNS | 80508 |  |  | *Thamnophilus* | *aethiops* | -5.2889 | -59.6936 |
| LSUMNS | 80716 |  |  | *Thamnophilus* | *aethiops* | -5.2518 | -59.6947 |
| LSUMNS | 81278 |  |  | *Thamnophilus* | *aethiops* | -4.0508 | -59.1153 |
| LSUMNS | 81338 |  |  | *Thamnophilus* | *aethiops* | -4.0508 | -59.1153 |
| LSUMNS | 86147 |  |  | *Thamnophilus* | *aethiops* | -6.7547 | -59.0831 |
| LSUMNS | 86229 |  |  | *Thamnophilus* | *aethiops* | -6.7517 | -59.0756 |
| LSUMNS | 86569 |  |  | *Thamnophilus* | *aethiops* | -6.5622 | -59.0939 |
| AMNH | J598 | MAR131 | 261259 | *Thamnophilus* | *aethiops* | -7.7037 | -60.5957 |
| AMNH | J616 | GT141 | 261282 | *Thamnophilus* | *aethiops* | -7.7037 | -60.5957 |
| LSUMNS | 85274 |  |  | *Thamnophilus* | *aethiops* | -5.7979 | -59.2308 |
| INPA | A15069 |  |  | *Thamnophilus* | *aethiops* | -3.3167 | -55.3333 |
| INPA | A16041 |  |  | *Thamnophilus* | *aethiops* | -4.5000 | -56.3000 |
| MPEG | T10229 | FPR085 |  | *Thamnophilus* | *aethiops* | -3.9461 | -58.4561 |
| MPEG | T14601 | ARA075 |  | *Thamnophilus* | *aethiops* | -2.7833 | -55.6000 |
| MPEG | T16625 | ARAII029 |  | *Thamnophilus* | *aethiops* | -3.0942 | -55.5356 |
| MPEG | T16704 | ARAII102 |  | *Thamnophilus* | *aethiops* | -2.5868 | -55.1952 |
| MPEG | T19436 | JAT(C)269 |  | *Thamnophilus* | *aethiops* | -5.0731 | -56.8559 |
| MPEG | T24591 | LCA081 |  | *Thamnophilus* | *aethiops* | -4.6167 | -56.6000 |
| MPEG | T742 | JRT060 |  | *Thamnophilus* | *aethiops* | -2.6000 | -56.1833 |
| MPEG | T7941 | JRT138 |  | *Thamnophilus* | *aethiops* | -2.4667 | -56.0000 |
| LSUMNS | 28095 |  |  | *Thamnophilus* | *aethiops* | -7.1333 | -75.6833 |
| LSUMNS | 75520 |  |  | *Thamnophilus* | *aethiops* | -10.3800 | -73.7200 |
| INPA | A7968 |  |  | *Thamnophilus* | *aethiops* | -4.9833 | -62.9667 |
| INPA | A8087 |  |  | *Thamnophilus* | *aethiops* | -4.9833 | -62.9667 |
| MPEG | T14205 | AMA341 |  | *Thamnophilus* | *aethiops* | -4.5303 | -71.6163 |
| INPA | A101 |  |  | *Willisornis* | *poecilinotus* | -7.2177 | -64.1827 |
| INPA | A10701 |  |  | *Willisornis* | *poecilinotus* | -4.8167 | -61.0000 |
| INPA | A1381 |  |  | *Willisornis* | *poecilinotus* | -4.1500 | -60.1333 |
| INPA | A265 |  |  | *Willisornis* | *poecilinotus* | -9.5285 | -65.3007 |
| INPA | A2725 |  |  | *Willisornis* | *poecilinotus* | -3.6833 | -60.3167 |
| INPA | A2749 |  |  | *Willisornis* | *poecilinotus* | -4.4167 | -60.9167 |
| INPA | A2759 |  |  | *Willisornis* | *poecilinotus* | -4.5833 | -61.2500 |
| INPA | A424 |  |  | *Willisornis* | *poecilinotus* | -5.1500 | -60.7333 |
| INPA | A7736 |  |  | *Willisornis* | *poecilinotus* | -4.9833 | -61.5667 |
| INPA | A9697 |  |  | *Willisornis* | *poecilinotus* | -5.2667 | -61.9167 |
| MPEG | T12400 | TUP066 |  | *Willisornis* | *poecilinotus* | -4.0833 | -60.6605 |
| MPEG | T13148 | OM096 |  | *Willisornis* | *poecilinotus* | -7.5183 | -63.3378 |
| LSUMNS | 25484 |  |  | *Willisornis* | *poecilinotus* | -2.7333 | -54.9153 |
| INPA | A12284 |  |  | *Willisornis* | *poecilinotus* | -4.7578 | -56.3925 |
| INPA | A14569 |  |  | *Willisornis* | *poecilinotus* | -5.2444 | -56.9175 |
| INPA | A16227 |  |  | *Willisornis* | *poecilinotus* | -4.7384 | -56.6217 |
| INPA | A7346 |  |  | *Willisornis* | *poecilinotus* | -5.0833 | -56.4333 |
| MPEG | T10694 | PIME138 |  | *Willisornis* | *poecilinotus* | -3.4729 | -54.5655 |
| MPEG | T12793 | PIME319 |  | *Willisornis* | *poecilinotus* | -6.5950 | -56.1015 |
| MPEG | T7132 | BMP050 |  | *Willisornis* | *poecilinotus* | -3.5271 | -52.3733 |
| MPEG | T9346 | AT011 |  | *Willisornis* | *poecilinotus* | -6.8427 | -56.1455 |
| INPA | A308 |  |  | *Willisornis* | *poecilinotus* | -8.8305 | -63.9605 |
| INPA | A3264 |  |  | *Willisornis* | *poecilinotus* | -9.2668 | -64.4001 |
| INPA | A334 |  |  | *Willisornis* | *poecilinotus* | -8.8533 | -63.8630 |
| INPA | A368 |  |  | *Willisornis* | *poecilinotus* | -8.9694 | -64.0699 |
| MPEG | T15888 | MAD064 |  | *Willisornis* | *poecilinotus* | -8.2239 | -63.1818 |
| MPEG | T15889 | MAD065 |  | *Willisornis* | *poecilinotus* | -8.2239 | -63.1818 |
| MPEG | T7625 | OP031 |  | *Willisornis* | *poecilinotus* | -10.8333 | -64.7500 |
| MPEG | T7627 | OP049 |  | *Willisornis* | *poecilinotus* | -10.8333 | -64.7500 |
| INPA | A24413 |  |  | *Willisornis* | *poecilinotus* | -8.4569 | -61.6427 |
| INPA | A4235 |  |  | *Willisornis* | *poecilinotus* | -7.7167 | -61.0667 |
| INPA | A465 |  |  | *Willisornis* | *poecilinotus* | -6.3000 | -60.4000 |
| INPA | A472 |  |  | *Willisornis* | *poecilinotus* | -6.3000 | -60.4000 |
| INPA | A882 |  |  | *Willisornis* | *poecilinotus* | -6.2500 | -60.4500 |
| INPA | A894 |  |  | *Willisornis* | *poecilinotus* | -7.8294 | -60.3963 |
| INPA | A896 |  |  | *Willisornis* | *poecilinotus* | -7.8294 | -60.3963 |
| INPA | A898 |  |  | *Willisornis* | *poecilinotus* | -7.8294 | -60.3963 |
| INPA | A902 |  |  | *Willisornis* | *poecilinotus* | -7.7322 | -60.9358 |
| MPEG | J163 | GDR210 |  | *Willisornis* | *poecilinotus* | -7.5672 | -60.6882 |
| MPEG | J192 | LJM199 |  | *Willisornis* | *poecilinotus* | -7.5672 | -60.6882 |
| MPEG | J202 | GT013 |  | *Willisornis* | *poecilinotus* | -7.5672 | -60.6882 |
| MPEG | J209 | GT016 |  | *Willisornis* | *poecilinotus* | -7.5672 | -60.6882 |
| MPEG | J222 | LJM207 |  | *Willisornis* | *poecilinotus* | -7.5672 | -60.6882 |
| MPEG | J226 | LJM208 |  | *Willisornis* | *poecilinotus* | -7.5672 | -60.6882 |
| MPEG | J302 | LJM221 |  | *Willisornis* | *poecilinotus* | -7.5672 | -60.6882 |
| MPEG | J439 | GDR289 |  | *Willisornis* | *poecilinotus* | -7.6951 | -60.9124 |
| MPEG | T2190 | MPDS698 |  | *Willisornis* | *poecilinotus* | -7.5500 | -62.5500 |
| MPEG | T3157 | MPDS752 |  | *Willisornis* | *poecilinotus* | -6.5500 | -62.0500 |
| MPEG | T388 | MAR084 |  | *Willisornis* | *poecilinotus* | -8.6957 | -61.4082 |
| MPEG | J335 | GDR256 |  | *Willisornis* | *poecilinotus* | -7.6998 | -60.8905 |
| MPEG | J340 | GDR257 |  | *Willisornis* | *poecilinotus* | -7.6998 | -60.8905 |
| MPEG | J391 | LJM240 |  | *Willisornis* | *poecilinotus* | -7.6998 | -60.8905 |
| MPEG | J421 | GDR284 |  | *Willisornis* | *poecilinotus* | -7.6998 | -60.8905 |
| AMNH | J644 | MAR137 | 261302 | *Willisornis* | *poecilinotus* | -8.1776 | -60.4531 |
| AMNH | J766 | MAR150 | 261448 | *Willisornis* | *poecilinotus* | -8.1776 | -60.4531 |
| AMNH | J768 | LJM323 | 261456 | *Willisornis* | *poecilinotus* | -8.1776 | -60.4531 |
| LSUMNS | 80798 |  |  | *Willisornis* | *poecilinotus* | -5.2518 | -59.6947 |
| LSUMNS | 81036 |  |  | *Willisornis* | *poecilinotus* | -5.7991 | -59.2686 |
| LSUMNS | 85474 |  |  | *Willisornis* | *poecilinotus* | -5.9333 | -59.1761 |
| LSUMNS | 86085 |  |  | *Willisornis* | *poecilinotus* | -6.7547 | -59.0831 |
| INPA | A891 |  |  | *Willisornis* | *poecilinotus* | -7.6373 | -60.6616 |
| AMNH | J561 | GDR329 | 261232 | *Willisornis* | *poecilinotus* | -7.7037 | -60.5957 |
| AMNH | J563 | LJM284 | 261226 | *Willisornis* | *poecilinotus* | -7.7037 | -60.5957 |
| AMNH | J610 | GDR348 | 261275 | *Willisornis* | *poecilinotus* | -7.7037 | -60.5957 |
| LSUMNS | 81265 |  |  | *Willisornis* | *poecilinotus* | -6.1633 | -59.0681 |
| INPA | A16298 |  |  | *Willisornis* | *poecilinotus* | -4.4950 | -56.3003 |
| MPEG | T10153 | FPR016 |  | *Willisornis* | *poecilinotus* | -4.0271 | -58.4349 |
| MPEG | T10234 | FPR100 |  | *Willisornis* | *poecilinotus* | -4.9144 | -58.2947 |
| MPEG | T11887 | PIME180 |  | *Willisornis* | *poecilinotus* | -5.8976 | -57.6966 |
| INPA | A815 |  |  | *Willisornis* | *poecilinotus* | -3.7500 | -66.0833 |
| MPEG | T23485 | JUT310 |  | *Willisornis* | *poecilinotus* | -3.2233 | -67.4417 |
| MPEG | T3645 | UFAC312 |  | *Willisornis* | *poecilinotus* | -9.7799 | -67.2186 |
| MPEG | T4097 | RUR022 |  | *Willisornis* | *poecilinotus* | -4.8667 | -65.1167 |

**Table S1:** List of all samples used in this study.

|  | ***Galbula*** | | | ***Malacoptila*** | | | ***Hypocnemis*** | | |
| --- | --- | --- | --- | --- | --- | --- | --- | --- | --- |
| **Population** | **π** | **θW** | **D** | **π** | **θW** | **D** | **π** | **θW** | **D** |
| West Inambari | 306.5 | 310.9 | -0.1 | 400.0 | 411.7 | -0.2 | - | - | - |
| East Inambari | 255.3 | 268.4 | -0.3 | 562.5 | 604.8 | -0.5 | - | - | - |
| Rondonia 1 | 281.3 | 281.3 | - | 427.3 | 427.3 | - | 111.3 | 111.3 | - |
| Rondonia 2 | 226.5 | 239.1 | -0.3 | 513.2 | 513.3 | 0.0 | 65.0 | 77.8 | -0.8 |
| Rondonia 2b | - | - | - | 564.2 | 578.7 | -0.3 | - | - | - |
| Rondonia 3 | 246.0 | 246.0 | - | 538.0 | 538.0 | - | 88.2 | 105.2 | -0.8 |
| Rondonia 4 | 242.5 | 265.7 | -0.5 | 568.0 | 568.0 | - | 99.8 | 142.0 | -1.3 |
| Rondonia 5 | 242.2 | 273.7 | -0.6 | 559.4 | 758.3 | -1.2 | 95.2 | 114.5 | -0.8 |
| Para | 185.7 | 228.4 | -0.9 | 501.5 | 629.2 | -1.0 | 94.7 | 110.2 | -0.8 |
|  | ***Thamnophilus*** | | | ***Phlegopsis*** | | | ***Willisornis*** | | |
| **Population** | **π** | **θW** | **D** | **π** | **θW** | **D** | **π** | **θW** | **D** |
| West Inambari | 22.6 | 25.0 | -0.7 | 1107.5 | 1118.7 | -0.1 | 400.2 | 417.8 | -0.4 |
| East Inambari | 25.3 | 32.8 | -1.1 | 1147.7 | 1310.6 | -0.6 | 310.4 | 409.3 | -1.1 |
| Rondonia 1 | 29.2 | 32.2 | -0.7 | 562.3 | 568.9 | -0.1 | 206.7 | 217.9 | -0.3 |
| Rondonia 2 | 23.3 | 30.7 | -1.2 | 604.7 | 651.0 | -0.3 | 245.2 | 340.8 | -1.2 |
| Rondonia 3 | 17.8 | 18.0 | -0.1 | 563.6 | 569.5 | -0.1 | 318.8 | 380.8 | -1.0 |
| Rondonia 4 | 20.0 | 27.6 | -1.4 | 645.4 | 676.3 | -0.2 | 252.0 | 284.6 | -0.6 |
| Rondonia 5 | 19.1 | 29.7 | -1.7 | 633.2 | 673.7 | -0.3 | 291.2 | 307.2 | -0.4 |
| Para | 27.2 | 28.9 | -0.4 | 696.8 | 772.3 | -0.5 | 225.2 | 270.1 | -0.9 |

**Table S2:** Population genetic statistics for each population within each species. Shown are nucleotide diversity (π), Watterson’s theta (θw), and Tajima’s D (Taj. D).

**Supplemental Figures:**

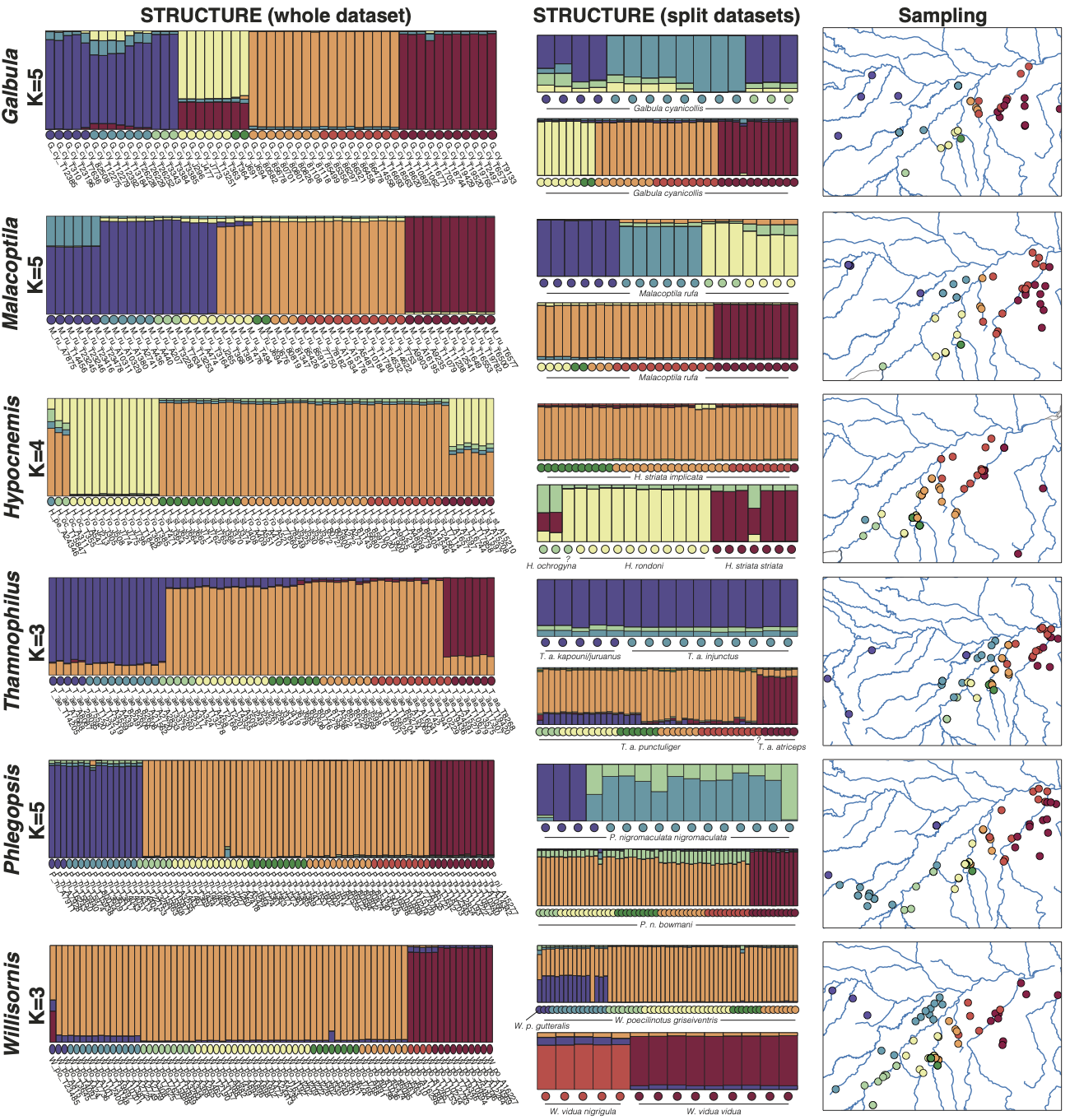

**Figure S1:** Results of best k-value STRUCTURE plots for the six species-groups. Structure plots in the left column show results with all individuals combined. The center column shows results for split groups. The map to the right of each species’ plot shows sampling locations. Circles under each STRUCTURE bar and on each map are colored by the *a priori* interfluve assignments (Figure 1 Map 1).

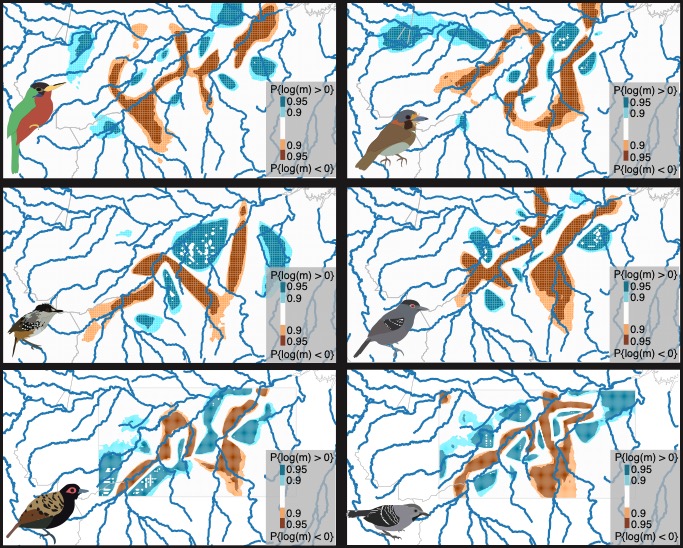

**Figure S2:** EEMS posterior probability of effective migration (m) estimated in EEMS for (left to right, top to bottom) *G. cyannicolis, M. rufa, Hypocnemis sp., T. aethiops, P. nigromaculata,* and *Willisornis sp*. Darker blues correspond to higher posterior probability of effective migration rate >0, whereas darker oranges correspond to higher posterior probability of effective diversity <0. Areas in white represent regions where the posterior probability log(m)≠0 is less than 0.9.

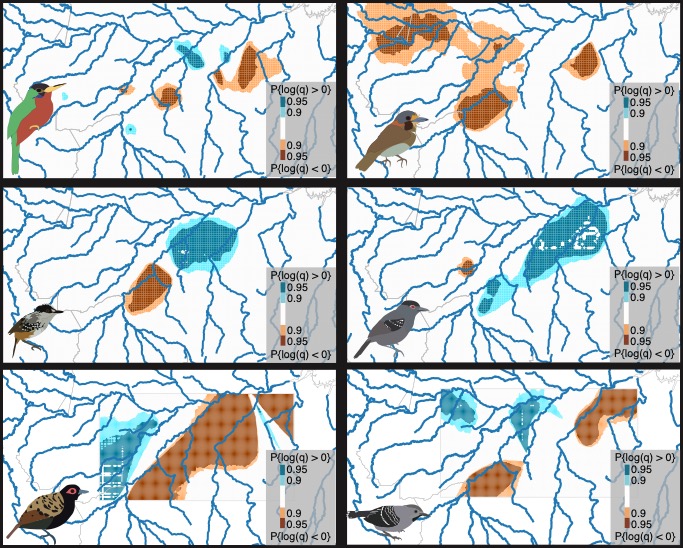

**Figure S3:** Posterior probability of effective diversity (q) estimated in EEMS for (left to right, top to bottom) *G. cyannicolis, M. rufa, Hypocnemis sp., T. aethiops, P. nigromaculata,* and *Willisornis sp*. Darker blues correspond to higher posterior probability of effective diversity >0, whereas darker oranges correspond to higher posterior probability of effective diversity <0. Areas in white represent regions where the posterior probability log(q)≠0 is less than 0.9.

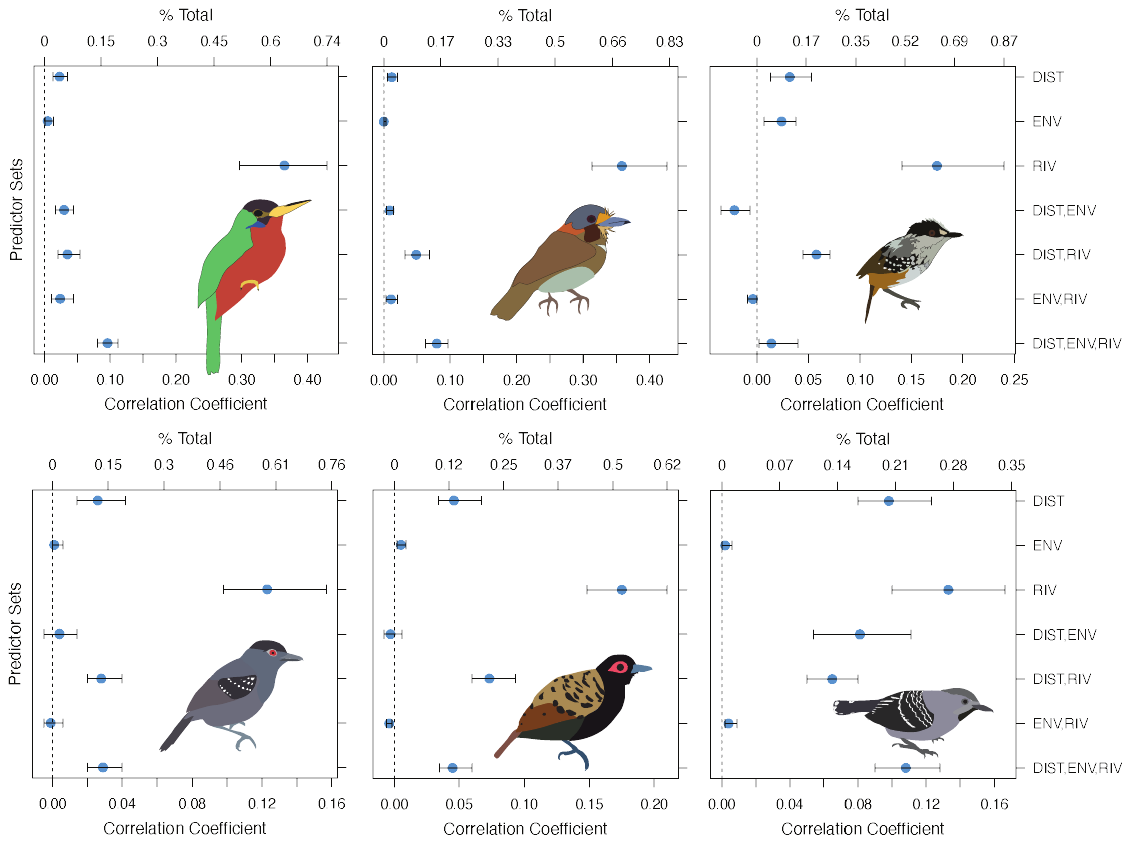

**Figure S4:** Results of commonality analysis. Rivers are the most important predictor of genomic divergence in five of the six species-groups based on a variance partitioning of the multivariate MMRR model, D ~ DIST + ENV + RIV, where D is genomic divergence in t-SNE space. The commonality coefficients of seven predictors (three unique and four common) for each species represent the proportion of variance of pairwise genomic divergence among samples that are explained by each predictor or set of predictors. The percent total represents the proportion of variance explained by the overall multivariate model that is explained by each predictor or set of predictors. Confidence intervals around each value were computed using 1,000 bootstrap replicates. The order of species is (from left to right beginning at the top) *Galbula, Malacoptila, Hypocnemis, Thamnophilus, Phlegopsis,* and *Willisornis*). Exact values for these parameters can be obtained in Table 2.

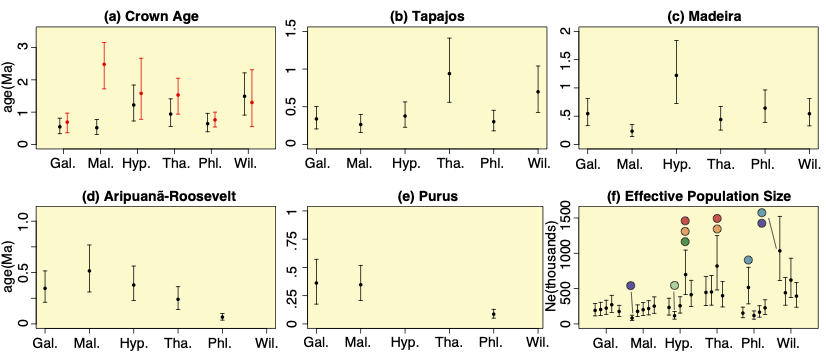

**Figure S5:** Parameter estimates from G-PhoCS demographic modeling. (a) Mean root (the basal split of all sampled populations) divergence times of each species-group inferred from G-PhoCS (black points) and 95% confidence intervals (black whiskers). The robustness of these results was assessed against data for the same divergence events as inferred from sanger sequencing data in a previous study (red points and whiskers) (*8*). (b–e) Divergence times across barriers showing mean values (black points) and 95% confidence intervals (whiskers), based on the allowed variance in mutation rate. (f) Estimates of mean (black points) effective population size (Ne) for each population of each species and 95% credible intervals assuming a fixed mutation rate. Populations within each species are ordered from southwest to northeast, and populations match those in PhyloNet and TreeMix. Colored circles indicating population assignments are shown for high or low values in each species.

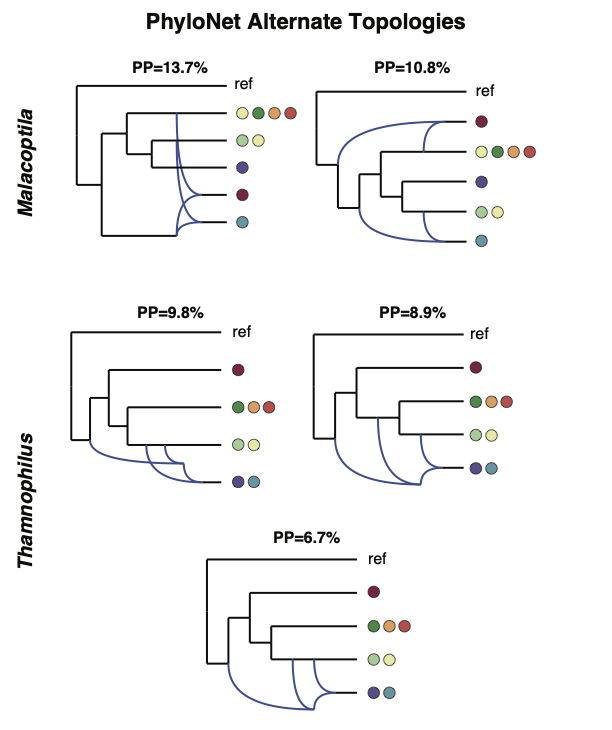

**Figure S6:** Alternate topologies from the PhyloNet analyses. The top two networks depict alternate topologies for *Malacoptila,* whereas the bottom three networks depict alternate topologies for *Thamnophilus*. These topologies differ very little from the most credible network topologies shown in Figure 5 of the main text.
